# Ancestral chromosomes for the Peronosporaceae inferred from a telomere-to-telomere genome assembly of *Peronospora effusa*

**DOI:** 10.1101/2021.09.14.460278

**Authors:** Kyle Fletcher, Oon-Ha Shin, Kelley J. Clark, Chunda Feng, Alexander I. Putman, James C. Correll, Steven J. Klosterman, Allen Van Deynze, Richard Michelmore

**Author notes:** Corresponding authors: Richard Michelmore, Allen Van Deynze.

## Abstract

We report the first telomere-to-telomere genome assembly for an oomycete. This assembly has extensive synteny with less complete genome assemblies of other oomycetes and will therefore serve as a reference genome for this taxon. Downy mildew disease of spinach, caused by the oomycete *Peronospora effusa*, causes major losses to spinach production. The 17 chromosomes of *P. effusa* were assembled telomere-to-telomere using Pacific Biosciences High Fidelity reads. Sixteen chromosomes are complete and gapless; Chromosome 15 contains one gap bridging the nucleolus organizer region. Putative centromeres were identified on all chromosomes. This new assembly enables a re-evaluation of the genomic composition of *Peronospora* spp.; the assembly was almost double the size and contained more repeat sequences than previously reported for any *Peronospora* spp. Genome fragments consistently under-represented in six previously reported assemblies of *P. effusa* typically encoded repeats. Some genes annotated as encoding effectors were organized into multigene clusters on several chromosomes. At least two effector-encoding genes were annotated on every chromosome. The intergenic distances between annotated genes were consistent with the two-speed genome hypothesis, with some effectors located in gene-sparse regions. The near-gapless assembly revealed apparent horizontal gene transfer from Ascomycete fungi. Gene order was highly conserved between *P. effusa* and the genetically oriented assembly of the oomycete *Bremia lactucae*. High levels of synteny were also detected with *Phytophthora sojae*. Many oomycete species may have similar chromosome organization; therefore, this genome assembly provides the foundation for genomic analyses of diverse oomycetes.

## Introduction

The Peronosporaceae are a family in the Oomycota with hundreds of described species, including *Phytophthora* and *Peronospora* spp. (Thines and Choi, 2016). Many species in the Peronosporaceae are downy mildews, which are obligate biotrophs that are typically host-specific plant pathogens. Other members of the Peronosporaceae are hemi-biotrophic, paraphyletic *Phytophthora* species that typically have a wider host range than downy mildew species (Abad et al., 2019). Phylogenetic analyses indicated that adaptation to obligate biotrophy from hemi-biotrophic ancestors has occurred at least twice within the family, such that downy mildews are polyphyletic (Bourret et al., 2018; Fletcher et al., 2018; Fletcher et al., 2019). *Peronospora* is the largest downy mildew genus and contains over 400 species (Thines and Choi, 2016), including *P. effusa*, the subject of the current analysis.

Downy mildew, caused by *P. effusa*, threatens spinach production because infection results in leaves unsuitable for sale and consumption. The disease is favoured by cool temperatures and high humidity and manifests as yellow lesions with blue-grey sporulation often on the abaxial surface of spinach leaves (Klosterman, 2016; Kandel et al., 2018). In conventional spinach production, the disease can be managed with use of fungicides and resistant cultivars; however, for organic production, deployment of resistant spinach cultivars is the only effective control available. Multiple resistance genes have been identified in the host, but the pathogen can rapidly adapt and evade host detection (Correll et al., 2011; Kandel et al., 2018). To date, there are 19 named races of *P. effusa;* however, isolates with variation in virulence phenotype are continuously identified and the biology behind the emergence of new races is not clearly understood (Plantum, 2021, April 15). Previous genomic investigations have yielded fragmented and repeat-sparse genome assemblies (Feng et al., 2018a; Fletcher et al., 2018; Klein et al., 2020). The number of chromosomes the genome of *P. effusa* contained, the number and genomic distribution of genes encoding virulence factors, and how much of the genome is conserved or variable between isolates remained unknown. Genome assemblies are important resources for determining the molecular basis for changes in virulence, resulting in the emergence of new pathotypes and provide informative loci for population studies.

In the current study, a 17-chromosome assembly is described for *P. effusa* that was generated using Pacific Biosciences High Fidelity (PacBio HiFi) reads. Sixteen telomere-to-telomere (T2T) gapless contigs and one T2T scaffold with a gap spanning the nucleolus organizer region were assembled. The genome was larger and more repeat-dense than anticipated and several new genes were annotated relative to previous assemblies. The content absent from previous assemblies was mostly long terminal repeat retrotransposons (LTR-RTs). There was a high degree of synteny between *P. effusa, P. sojae*, and *Bremia lactucae*. Putative centromeres were identified through comparative genomics with *P. sojae*. This is a landmark genome assembly for oomycete genomics.

## Results

To determine the virulence phenotype of isolate UA202013, disease incidence was recorded on each of the standardized differential spinach lines. Disease incidence indicated that isolate UA202013 had a virulence pattern (Table 1), which did not match any of the 19 denominated races of *P. effusa* (Plantum, 2021, April 15).

**Table 1.** Disease incidence and reactions of spinach differential cultivars to isolate UA202013 of *P. effusa*.

| Differential Line | Disease Incidence (%) <sup>a</sup> | Disease Reactions <sup>b</sup> |
| --- | --- | --- |
| Viroflay | 100.0 | + |
| NIL5 | 26.7 – 70.0 | +/- |
| NIL3 | 100.0 | + |
| NIL4 | 15.4 – 64.3 | +/- |
| NIL6 | 100.0 | + |
| NIL1 | 0.0 | - |
| NIL2 | 0.0 | - |
| Pigeon | 0.0 | - |
| Caladonia | 0.0 | - |
| Meerkat | 0.0 | - |
| Hydrus | 0.0 | - |

<sup>a</sup> Disease incidence is based on the number of plants (cotyledon or true leaves) showing chlorosis and/or sporulation out of the total plants evaluated
<sup>b</sup> Differential lines were considered susceptible (+) if disease incidence was greater than 85% or resistant (-) if disease incidence was less than 15%. The disease reaction was considered intermediate (+/-) if disease incidence was between 15–85%.

The genome of isolate UA202013 of *Peronospora effusa* was assembled into 17 telomere-to-telomere chromosomes. Sixteen of the chromosomes are complete and gapless; Chromosome (Chr.) 15 contains a single gap bridging the nucleolus organizer region (NOR). Chromosome assembly length ranged from 1.78 Mb (Chr. 9) to 8.42 Mb (Chr. 1; Fig. 1). Putative centromeres were identified on all 17 chromosomes (Fig. 1); 15 putative centromeres contained at least one Copia-like transposon (CoLT) element (Supplementary Fig. 1). CoLT elements were previously hypothesized to be enriched in oomycete centromeres (Fang et al., 2020). Twelve *P. effusa* centromeres were syntenic with experimentally validated *P. sojae* centromeres (Fang et al., 2020), though some gene rearrangements were present around the centromeres of *P. effusa* Chromosomes 1, 6, 7, 8, and 15 (Supplementary Fig. 1). Putative centromeres of *P. effusa* Chr. 1 and Chr. 15 did not contain CoLTs but were syntenic with *P. sojae* centromeres (Supplementary Fig. 1). Telomeric satellite sequences (5’-CCCTAAA-3’) were detected at both ends of every chromosome and ranged from 321 bp to 1,283 bp. The GC content of the *P. effusa* assembly was 48.6%, ranging from 49.2% to 48.1% per chromosome (Table 3). Within 100 kb windows, the AT content ranged from 60.9% to 43.1% (Fig. 1A). The assembly was scored by BUSCO (Simao et al., 2015) as 99.2% complete (232/234 models), of which 0.9% (2 models) were duplicated, using a protist-specific database.

**Fig. 1.**
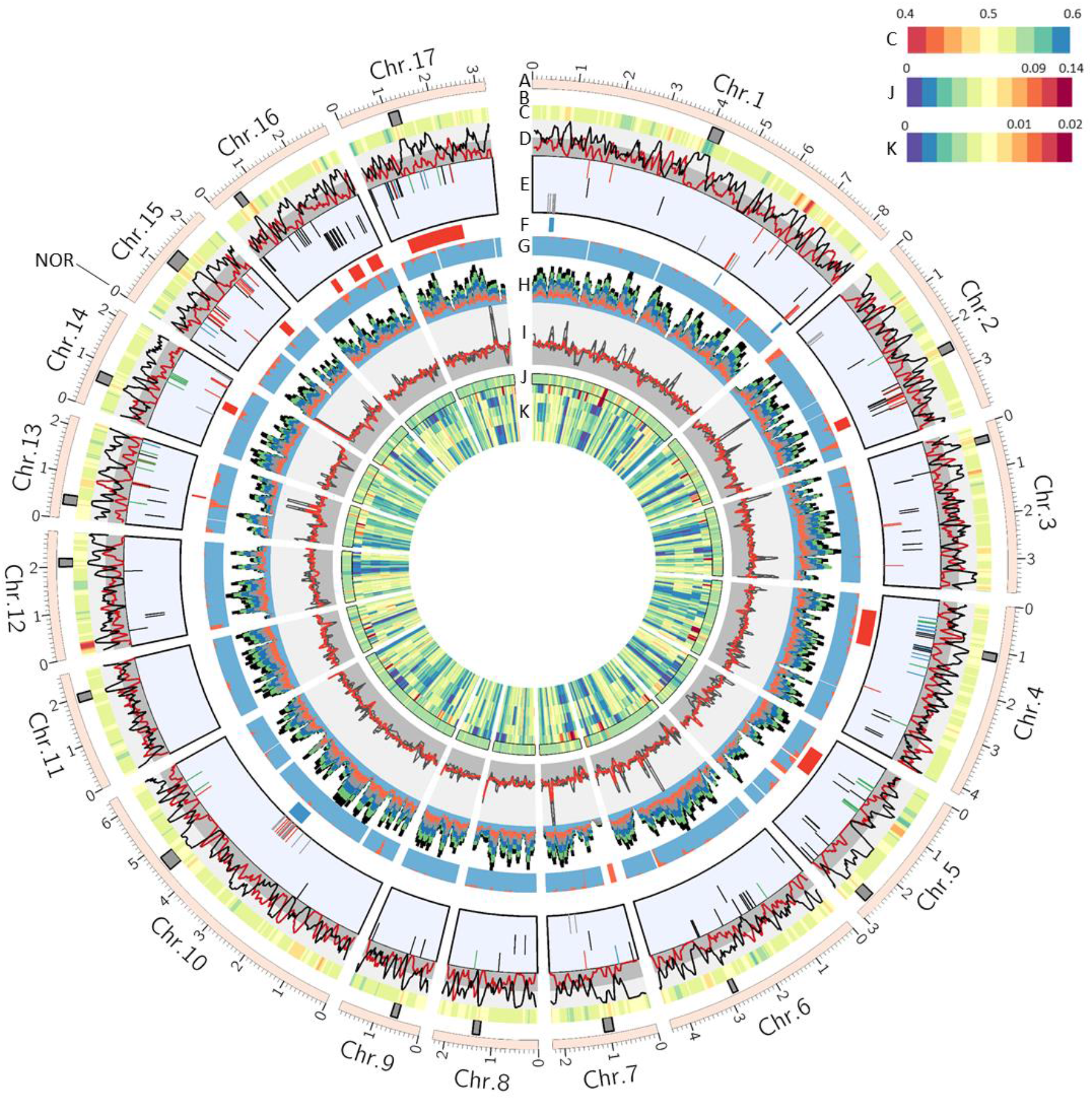
Architecture of the 17 chromosomes of *Peronospora effusa*. A) Scaled chromosomes and ticks showing lengths. Chromosome number was designated based on syntenic linkage groups with *B. lactucae*. B) Dark grey boxes indicate putative positions of centromeres, some of which are syntenic with *P. sojae* (Supplementary Fig. 1). All subsequent tracks are plotted in 100 kb bins, with a 25 kb step. C) Heatmap of AT content, ranging from 0.4 to 0.6. D) Gene density depicted in red, repeat density depicted in black. The dark grey background shows 0 to 0.5, light grey 0.5 to 1. E) Effector annotations. The first row are WY effectors; red ticks indicate the annotation contains a signal peptide prediction, RXLR and EER motif and WY domain; blue ticks indicate annotation with a signal peptide and RXLR motif and WY domain; green ticks indicate annotation with signal peptide and WY domain; black ticks indicate a WY domain annotation with no signal peptide. In the second row are genes annotated as encoding RXLR-EER proteins with a signal peptide (all black). Third row are crinkler annotations with (red) and without (black) predicted signal peptides. F) Clusters of high identity crinkler (clue) and RXLR-EER/WY (red) annotation. G) Orthology detected with 33 other oomycetes, blue indicates ortholog detected with other species, red indicates genes unique to *P. effusa*, white indicates no annotation in a bin. H) Coverage of bin in six smaller *P. effusa* assemblies as determined by BLASTn. Colours indicate different assemblies; all are Illumina-based except the bottom light blue, which was a hybrid Pac Bio and Illumina assembly. Expanded in Supplementary Fig. 8. I) Normalized read depth of each bin, long reads of UA202013 (33x) were plotted in red. Short reads of five other isolates were plotted in black. The background indicated 0x to 1x (dark grey) and 1x to 3x (light grey). Expanded in Supplementary Fig. 14. J) Density of structural variant calls inferred by Freebayes using the long reads of UA202013, plotted with scale log base = 0.5. K) Single nucleotide variant calls from alignments against UA202013 for alignments of UA202013 long reads, and short reads of Pfs12, Pfs13, R13, Pfs14, and R14 plotted with scale log base = 0.5. Heterozygosity is plotted in Supplementary Fig. 18.

The genome was 53.7% repetitive. The most common repeat elements were long terminal-repeat retrotransposons (LTR-RTs), comprising 40.2% of the genome. Chromosomes ranged from 43.1% to 67.1% repetitive sequence. Repeat content did not correlate with chromosome length (Table 2; Pearson’s Correlation = -0.38). The total genome size of 58.6 Mb was approximately twice the size of previous assemblies (23.9 Mb to 32.1 Mb). Repeat content calculated for previous assemblies ranged from 14.4% to 39.0% (Table 3). Therefore, most of the size difference between the new assembly and previous assemblies is due to the repeat content assembled. Putative centromeres co-located with repetitive regions (Fig. 1). Pairwise alignment of 66,858 clustered repeat elements demonstrated that many repeats shared high identity (≥98%; Supplementary Fig. 2). High identity alignments of repeats were found both within and between chromosomes (Supplementary Fig. 3). Nucleotide differences between LTR pairs of retrotransposons in the new assembly were less diverged than in previous *P. effusa* assemblies (Supplementary Fig. 4). Therefore, similar LTR-RTs assembled in short read assemblies were likely chimeric, pairing LTRs from different elements. This suggests that short-read assemblies are inadequate to date LTR expansion events accurately in oomycetes. Interestingly, the LTR divergence in the new HiFi assembly was comparable to that reported for *P. ramorum* and *P. sojae*, both assemblies generated by Sanger sequencing (Tyler et al., 2006).

**Table 2.** Summary statistics of chromosomes assembled for *Peronospora effusa* isolate UA202013.

| *P. effusa*<br>UA202013 | Assembled<br>length (bp) | Total gaps<br>(bp) | Percent<br>repeats | AT content | Genes<br>annotated | Genes/Mb | Effectors<br>annotated | 5' Telomere<br>assembled<br>length (bp) | 3' Telomere<br>assembled<br>length (bp) |
| --- | --- | --- | --- | --- | --- | --- | --- | --- | --- |
| Chr. 1 | 8,417,896 | 0 | 51.5% | 51.4% | 1482 | 176 | 70 | 706 | 685 |
| Chr. 2 | 3,956,849 | 0 | 53.5% | 51.7% | 596 | 151 | 23 | 597 | 958 |
| Chr. 3 | 3,727,724 | 0 | 49.5% | 50.9% | 666 | 179 | 10 | 723 | 616 |
| Chr. 4 | 4,163,955 | 0 | 51.4% | 51.3% | 751 | 180 | 28 | 803 | 695 |
| Chr. 5 | 3,019,954 | 0 | 57.3% | 51.5% | 393 | 130 | 16 | 680 | 1039 |
| Chr. 6 | 4,409,438 | 0 | 54.9% | 51.5% | 692 | 157 | 14 | 1283 | 649 |
| Chr. 7 | 2,262,584 | 0 | 67.1% | 51.5% | 261 | 115 | 6 | 756 | 703 |
| Chr. 8 | 2,258,044 | 0 | 56.3% | 51.6% | 334 | 148 | 5 | 502 | 656 |
| Chr. 9 | 1,779,483 | 0 | 49.7% | 51.8% | 315 | 177 | 3 | 682 | 523 |
| Chr. 10 | 6,355,493 | 0 | 48.6% | 51.3% | 1185 | 186 | 26 | 669 | 652 |
| Chr. 11 | 2,566,028 | 0 | 43.1% | 51.8% | 563 | 219 | 0 | 707 | 867 |
| Chr. 12 | 2,778,632 | 0 | 54.8% | 50.8% | 577 | 208 | 4 | 785 | 505 |
| Chr. 13 | 2,196,953 | 0 | 63.0% | 51.2% | 237 | 108 | 17 | 603 | 321 |
| Chr. 14 | 2,153,540 | 0 | 61.1% | 51.6% | 310 | 144 | 12 | 801 | 619 |
| Chr. 15 | 2,369,557 | 100 | 62.3% | 51.8% | 281 | 119 | 24 | 525 | 654 |
| Chr. 16 | 2,944,237 | 0 | 54.1% | 51.3% | 479 | 163 | 27 | 795 | 784 |
| Chr. 17 | 3,218,066 | 0 | 52.8% | 51.9% | 623 | 194 | 22 | 708 | 553 |
| Total | 58,578,433 | 100 | 53.7% | 51.4% | 9745 | 166 | 307 |  |  |

**Table 3.** Comparative assembly statistics of *P. effusa* isolate UA202013 with previously published assemblies of *P. effusa*.

| Isolate | Scaffold N50 | Scaffold count | Contig N50 | Contig count | Assembly size | Gaps | Repeats <sup>1</sup> | Sequencing technologies | Reported gene models | Study |
| --- | --- | --- | --- | --- | --- | --- | --- | --- | --- | --- |
| UA202013 | 3.7 Mb | 17 | 3.7 Mb | 18 | 58.6 Mb | <0.001% | 53.7% | PacBio HiFi | 9,745 | This study |
| R13 | 72 kb | 784 | 48 kb | 1472 | 32.2 Mb | 0.26% | 34.5% | Illumina | 8,607 | (Fletcher et al., 2018) |
| R14 | 61 kb | 880 | 52 kb | 1275 | 30.8 Mb | 0.56% | 32.0% | Illumina | 8,571 | (Fletcher et al., 2018) |
| Pfs1 | 33.7 kb | 6762 | 33 kb | 6777 | 32.1 Mb | 0.02% | 39.0% | Illumina + PacBio | 13,277 <sup>2</sup> | (Klein et al., 2020) |
| Pfs12 | 21.3 kb | 4061 | 20.9 kb | 4505 | 25.2 Mb | 0.2% | 17.5% | Illumina | Not Reported | (Feng et al., 2018a) |
| Pfs13 | 18.1 kb | 4387 | 17.8 kb | 4939 | 24 Mb | 0.26% | 14.4% | Illumina | Not Reported | (Feng et al., 2018a) |
| Pfs14 | 22.1 kb | 4020 | 21.3 kb | 4488 | 24.9 Mb | 0.22% | 16.6% | Illumina | Not Reported | (Feng et al., 2018a) |

<sup>1</sup> Repeat content calculated in the present study using a repeat library calculated from *P. effusa* UA202013.
<sup>2</sup> Inflated gene model count due to some repeat elements annotated as genes.

A total of 9,745 genes were annotated in the *P. effusa* genome, with an average of 166 genes per Mb. The number of genes annotated per chromosome ranged from 237 to 1,482 (Table 2) and correlated with chromosome length (PC = 0.97). The density of genes ranged from 108 genes per Mb to 219 genes per Mb. Gene density did not correlate with chromosome length (PC = 0.29). On average, 27.6% of each 100 kb bin encoded genic sequence. Gene density was inversely related to repeat density (Fig. 1). Of these genes, 307 were annotated as effectors. These included 98 crinkler effectors and 209 RXLR effectors. Effectors were encoded on every chromosome except Chr. 11 (Table 2). The effector count per chromosome was weakly correlated with the chromosome length (PC = 0.82). Some effectors were organized as clusters of genes with high identity (Fig. 1, Supplementary Fig. 5). Three clusters of crinkler annotations on Chr. 1, spanning approximately 141 kb, 34 kb, and 84 kb, were detected. There was a fourth, larger crinkler cluster on Chr. 10, spanning 523 kb. Clusters of proteins annotated as having an RXLR-EER motif, WY domain, or both (RXLR-EER/WY) were identified on Chr. 2 (310 kb), Chr. 4 (1.1 Mb), Chr. 5 (745 kb), Chr. 14 (249 kb), Chr. 15 (245 kb), Chr. 17 (168 kb), and three independent clusters on Chr. 16 (182 kb, 360 kb, and 378 kb; Fig. 1; Supplementary Fig. 5). The RXLR-EER/WY clusters on Chr. 4 and Chr. 17 spanned centromeres that shared synteny with *P. sojae* (Supplementary Fig. 1). Inter-chromosome, lower identity alignments were detected for RXLR-EER/WY annotations, but not crinkler annotations (Supplementary Fig. 5). These inter-chromosomal alignments could be due to gene duplication and transposition. Crinkler clusters were composed of protein sequences sharing higher identity to one another than RXLR-EER/WY clusters (Supplementary Fig. 6). Inter-chromosomal alignments of effector proteins rarely shared high identity compared to intra-chromosomal alignments (Supplementary Fig. 6).

Inferring orthology with the annotation of 37 other oomycetes from 34 different species (Supplementary Table 1) assigned 8,739 annotations of *P. effusa* UA202013 to 6,425 orthogroups. An additional 725 UA202013 proteins assigned to 483 orthogroups were identified as unique to *P. effusa* and annotated in previous assemblies. Therefore, 281 *P. effusa* UA202013 annotations had no orthology assigned with any previous oomycete assembly. Genes unique to *P. effusa* could be found on every chromosome (Supplementary Fig. 7). Of the 1,006 genes unique to *P. effusa*, 174 were annotated as effectors and a further 54 overlapped effector clusters.

The complete T2T genome assembly allowed for the investigation of genes unique to *P. effusa* as potential horizontal gene transfer (HGT) events. An obvious signal for HGT was two neighbouring gene models in the sub-telomeric region of Chr. 2 of *P. effusa*. The translated sequence of these two gene models was highly homologous to several fungal proteins. The fungal homologs also neighboured one another in their respective genome assemblies (Fig. 2A). One of these genes encoded a metallophosphatase domain (MPD); the second gene did not encode a detectable conserved domain. Both were annotated in previous *P. effusa* assemblies (Fletcher et al., 2018). The genes were embedded in a repeat-rich region on Chr. 2, ∼22 kb and ∼64 kb away from flanking genes (Fig. 2A). Phylogenetically, the *P. effusa* genes nested within fungal protein sequences, closest to the sunflower pathogen *Diaporthe helianthi* and dieback associated fungi *Valsa sordida* and *V. malicola* (Fig. 2B, 2C). The flanking genes on *P. effusa* Chr. 2 were assigned orthology with other oomycete proteins and branched as expected phylogenetically (Fig. 2D, 2E). Flanking gene blocks were conserved between oomycete assemblies, but rearrangements of the gene blocks were apparent (Fig. 2A). A paralogous MPD encoding gene was present on *P. effusa* Chr. 15. There was no paralog for the second HGT candidate gene in this region. The region on Chr. 15 was flanked by a block of oomycete genes and five copies of a repeated sequence with homology to the flanking repeat on Chr. 2 (Supplementary Fig. 8). Additional MPD encoding homologous genes were annotated on *P. effusa* Chr. 1 and Chr. 13. The intergenic distance flanking these genes was less than 500 bp. These genes were assigned orthology with other oomycete MPD encoding genes and branched as expected phylogenetically (Fig. 2). Therefore, MPD genes on Chr. 1 and Chr. 13 were not acquired by HGT. Genes encoding an MPD domain on Chr. 2 and Chr. 15 were likely acquired through a single HGT event from true fungi, followed by duplication. The read-depth of these genes in R15 and five other previously sequenced isolates was consistent with all four genes being present in all isolates sequenced (Supplementary Table 2).

**Fig. 2.**
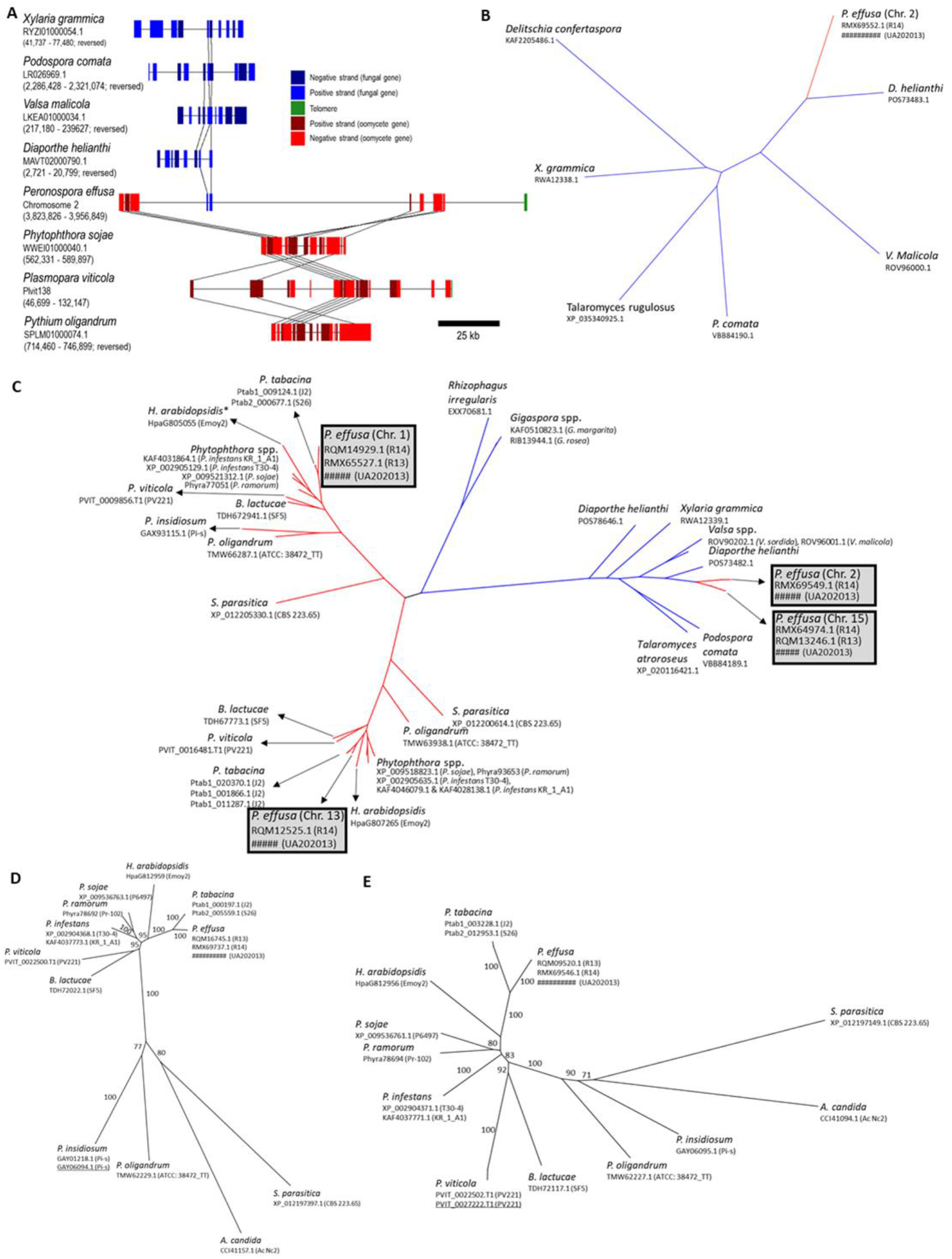
Evidence of horizontal gene transfer in the genome of *Peronospora effusa*. A) Gene content of several fungal and oomycete contigs or scaffolds. The top four sequences are true fungi and show conserved gene order of two to three genes, linked by black lines. Two of those genes were found on Chr. 2 of *P. effusa*, coloured blue and embedded in repeats. The red blocks indicate oomycete genes. Orthologs of oomycete genes were found on single contigs/scaffolds in the genomes of other oomycetes. The gene order of *P. sojae* and *P. oligandrum* was similar. Rearrangements were visible when the *P. sojae* and *P. oligandrum* sequences were compared to *P. effusa* or *P. viticola*. B–E) Neighbour joining trees of the horizontally acquired genes and the flanking gene. B) Only found in *P. effusa*, the protein sequence was nested within fungal sequences. C) Several genes encoding proteins containing metallophosphatase domains were found in the *P. effusa* genome. Homologs on Chr. 2 and Chr. 15 were nested in fungal sequences; no orthologs were annotated in oomycete assemblies. Orthologs in the genomes of other oomycetes were identified for the homologs on *P. effusa* Chr. 1 and Chr. 13. Phylogenetic analysis of the glancing proteins (D and E) produced expected topologies. Annotation of UA20213 submitted to NCBI and awaiting the assignment of accession identities. Similar protein sequences can be found by using the NCBI accessions of P. effusa isolates R13 or R14.

Alignment of six previous *P. effusa* genome assemblies confirmed that most sequence missing from these previous assemblies was repeat encoding and not genic. The percentage of each 100 kb window covered by BLASTn alignments generated with other *P. effusa* assemblies was calculated, revealing poor representation in multiple windows across all chromosomes (Fig. 1E; Supplementary Fig. 9). This result is consistent with the previous assemblies being smaller than the new draft assembly (Table 3). Between 1,140 genes and 1,655 genes annotated in *P. effusa* UA202013 were absent in the assemblies of other isolates based on BLASTn alignments. As previously noted, the inflated gene count of *P. effusa* isolate Pfs1 is likely due to repeats annotated as genes (Klein et al., 2020). In total, 3,620 genes annotated in *P. effusa* UA202013 were not covered by BLASTn alignments from at least one other assembly of *P. effusa*. Only 121 genes were inferred as absent from all other assemblies of *P. effusa* (Supplementary Fig. 10). Further investigation revealed that 98 of the 121 proteins lacked orthology with any oomycete protein and may therefore be unique to isolate UA202013 of *P. effusa* or annotation artefacts.

Orthology analysis of 38 annotated oomycete genomes, including three from *P. effusa*, supported little novelty in the UA202013 assembly compared to *P. effusa* isolates R13 and R14. Isolate UA202013 was assigned to 97 multi-species orthogroups that did not contain R13 or R14 genes. Genes of all three isolates were assigned to a total of 6,264 orthogroups (Supplementary Fig. 11). These results suggest that most of the gene space is accessible through short-read assembly and that there are very few unique gene annotations specific to isolate UA202013. Further analysis of orthologs showed that the increased gene count for UA202013 is due to collapsed paralogs in other assemblies. There were several instances of orthogroups being assigned multiple annotations in the UA202013 assembly but having fewer or no annotations in R13 or R14 assemblies (Supplementary Fig. 12). Analysis of read depth supported the correct resolution of paralogs in the UA202013 assembly; high coverage of genes in multiple orthogroups in R13 and R14 demonstrates that the paralogs were collapsed in the short-read assemblies (Supplementary Fig. 13). Therefore, short-read assemblies are still a valuable resource for obtaining the gene space of these organisms, though they are not sufficient to resolve the highly repetitive architecture nor sufficient to resolve paralogous genes in the genome of *P. effusa*.

Read-depth analysis showed no copy number variation of whole chromosomes was present in isolates UA202013, Pfs12, Pfs13, Pfs14, R13, and R14 (Fig. 1I; Supplementary Fig. 14). The only significant discrepancy in coverage was observed at the gap on Chr. 15 when aligning reads of UA202013 back to the UA202013 assembly. This high coverage sequence encoded ribosomal subunits, consistent with it being the NOR. This increased coverage across the NOR was detected in the reads of other isolates (Fig. 1I; Supplementary Fig. 14). Additional significant peaks and troughs of coverage were recorded for the five other isolates. Read depth differences correlated with the phenotypes, suggesting the genomes of Pfs13 and R13 were similar, as were the genomes of Pfs12, Pfs14, and R14 (Supplementary Fig. 14). Interestingly, few reads for all five other isolates aligned to the effector cluster on Chr. 14 of UA202013 (Supplementary Fig. 14, 15). This result suggests that the effectors encoded may be absent or highly diverged in UA202013 versus the other five isolates. The four effectors annotated in UA202013 contained a signal peptide prediction, an EER motif, two LWY domains, a partial GLY zipper domain, and a partial electron transport complex protein domain (Supplementary Fig. 16). Between all four, the N-terminus was more conserved than the C-terminus. The genomes of R13 and R14 each contained a single homologous annotation, though in both cases the partial electron transport complex protein domain was not assembled as in the UA202013 gene models (Supplementary Fig. 16). No significant depth variation was observed for other effector clusters in the UA202013 assembly (Supplementary Fig. 14). Therefore, the cluster of putative effectors on Chr. 14 contains candidates for proteins underlying the difference in virulence of isolate UA202013 compared to the other isolates.

Loss of heterozygosity (LOH) on Chr. 6 may underlie the difference in virulence between races 12 and 14 of *P. effusa*. Races 12 and 14 have an identical virulence phenotype except race 14 is virulent on the differential cultivar Pigeon and race 12 is not (Plantum, 2021, April 15). Comparisons of all six isolates in this analysis revealed that the race 14 isolates, R14 and Pfs14, and race 12 isolate, Pfs12, were more similar to one another genome-wide than to other isolates. Among these three isolates, clustering of SNPs consistently grouped Pfs14 with Pfs12, suggesting they were genetically closer to each other than either was to R14 (Supplementary Fig. 17). However, the two independently collected race 14 isolates (Feng et al., 2018b; Fletcher et al., 2018) were both highly homozygous compared to Pfs12 over two regions on Chr. 6 and Chr. 14 (Supplementary Fig. 18, 19). Therefore, LOH in one or both of these regions may have resulted in the loss of an avirulence allele detectable by cultivar Pigeon. In the assembly of UA202013, seven genes were annotated as encoding effectors in the 750 kb region on Chr. 6. No effector encoding genes were annotated in the Chr. 14 region (Supplementary Fig. 18, 19). Therefore, one or more of these genes are good candidates for determining the avirulence phenotype.

To establish if the genome of *P. effusa* was compartmentalized into gene dense and gene sparse regions, the distribution of intergenic distances between all genes was calculated. This revealed a bimodal distribution for both the 5’ and 3’ flanking regions (Fig. 3, Supplementary Fig. 20). Most genes (80.5%) were flanked by 100 bp to 5 kb of intergenic sequence on both sides. Approximately 17%, or 1,675 annotations had intergenic distances greater than 5 kb on one side, of which 221 had >5 kb of intergenic sequence flanking on both sides. This included 142 and 40 effector annotations, meaning 8.5% and 18% of the genes in these respective compartments were effectors (Fig. 3). Genome-wide, 3.1% of the genes are annotated as effectors; therefore, effectors were enriched in the gene-sparse regions of the assembly. The upper limit of this analysis was better than a previously reported similar analysis of *P. effusa* Pfs1 (Klein et al., 2020), likely because the Pfs1 assembly was more fragmented than the T2T UA202013 assembly. The two-speed genome hypothesis proposes that effectors are embedded in gene-sparse regions of the genome (Dong et al., 2015). Genes with >50 kb intergenic regions were found on every chromosome, suggesting that every chromosome of *P. effusa* has potentially flexible regions involved in pathogenicity.

**Fig. 3.**
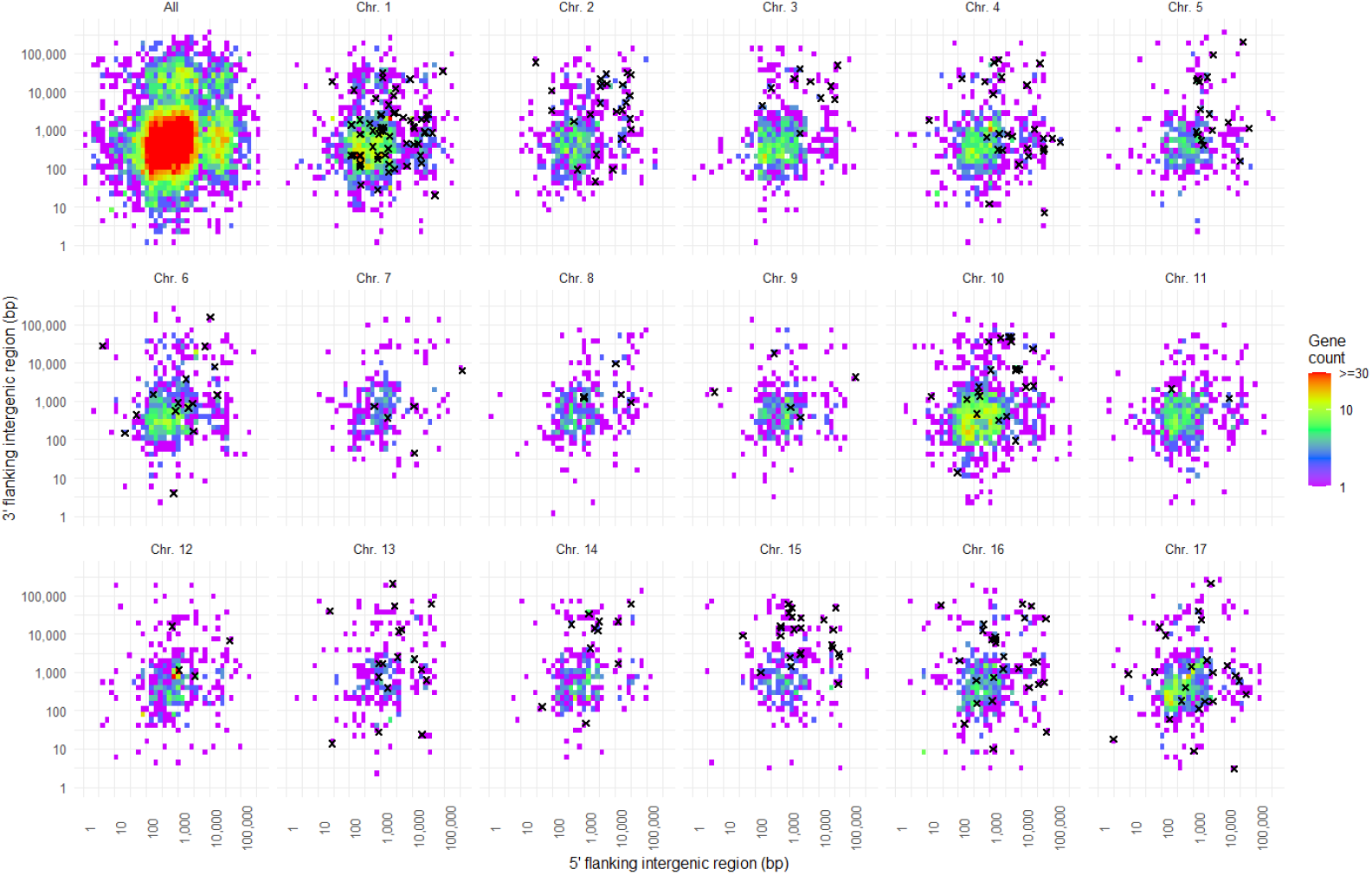
Intergenic distances between annotated genes of *Peronospora effusa* isolate UA202013. For each gene, the distance 5’ to the next gene are plotted on the x-axis and the distance 3’ to the next gene are plotted on the y-axis. The top left plot shows the intergenic distance between all annotations showing that the majority of genes cluster between 100 bp and 5,000 bp on either side (coloured red). Some genes had larger intergenic distances on either, or both sides. The other plots show individual chromosomes; black ‘x’s mark genes annotated as encoding effectors.

Comparative genomics demonstrated that the chromosomes of *P. effusa* were highly syntenic with assemblies of other oomycetes (Fig. 4A, 4B). The genome of *B. lactucae* has been ordered into 19 linkage groups (Fletcher et al., 2020), 15 of which were colinear with complete chromosomes of *P. effusa* (Fig. 4A). The other four *B. lactucae* linkage group (LG) scaffolds were syntenic with two chromosomes of *P. effusa*: LG 16 and LG 19 of *B. lactucae* were syntenic with Chr. 16 of *P. effusa*; LG 17 and LG 18 of *B. lactucae* were syntenic with Chr. 17 of *P. effusa*. Gene content was also conserved with *P. sojae* (Fig. 4B), though synteny of some scaffolds was found with multiple chromosomes (*P. sojae* 116 = *P. effusa* Chr. 1 and Chr. 13; *P. sojae* 115 = *P. effusa* Chr. 6 and Chr. 10, *P. sojae* 117 = *P. effusa* Chr. 9, Chr. 11, and Chr. 12). It remains unresolved whether these are true chromosomal fusions or assembly errors in *P. sojae*. Comparison to a more recent long-read assembly of *P. sojae* did not resolve this because the assembly was too fragmented (Supplementary Fig. 1). Phylogenetics of BUSCO genes found to be single copy across 31 oomycete assemblies supports polyphyly of downy mildews as reported previously (Bourret et al., 2018; Fletcher et al., 2018; Fletcher et al., 2019). The most recent common ancestor of *P. effusa* and *B. lactucae* is inferred as common to all downy mildews and *Phytophthora* species clades one to five (Fig. 4C). The most recent common ancestor of *P. effusa* and *P. sojae* (*Phytophthora* clade 7) is more ancient than the common ancestor of *P. effusa* and *B. lactucae*. Given the high levels of synteny between these three species, it is likely that the 17-chromosome architecture of *P. effusa* and *B. lactucae* will be ancestral to hundreds of other species in the Peronosporaceae.

**Fig. 4.**
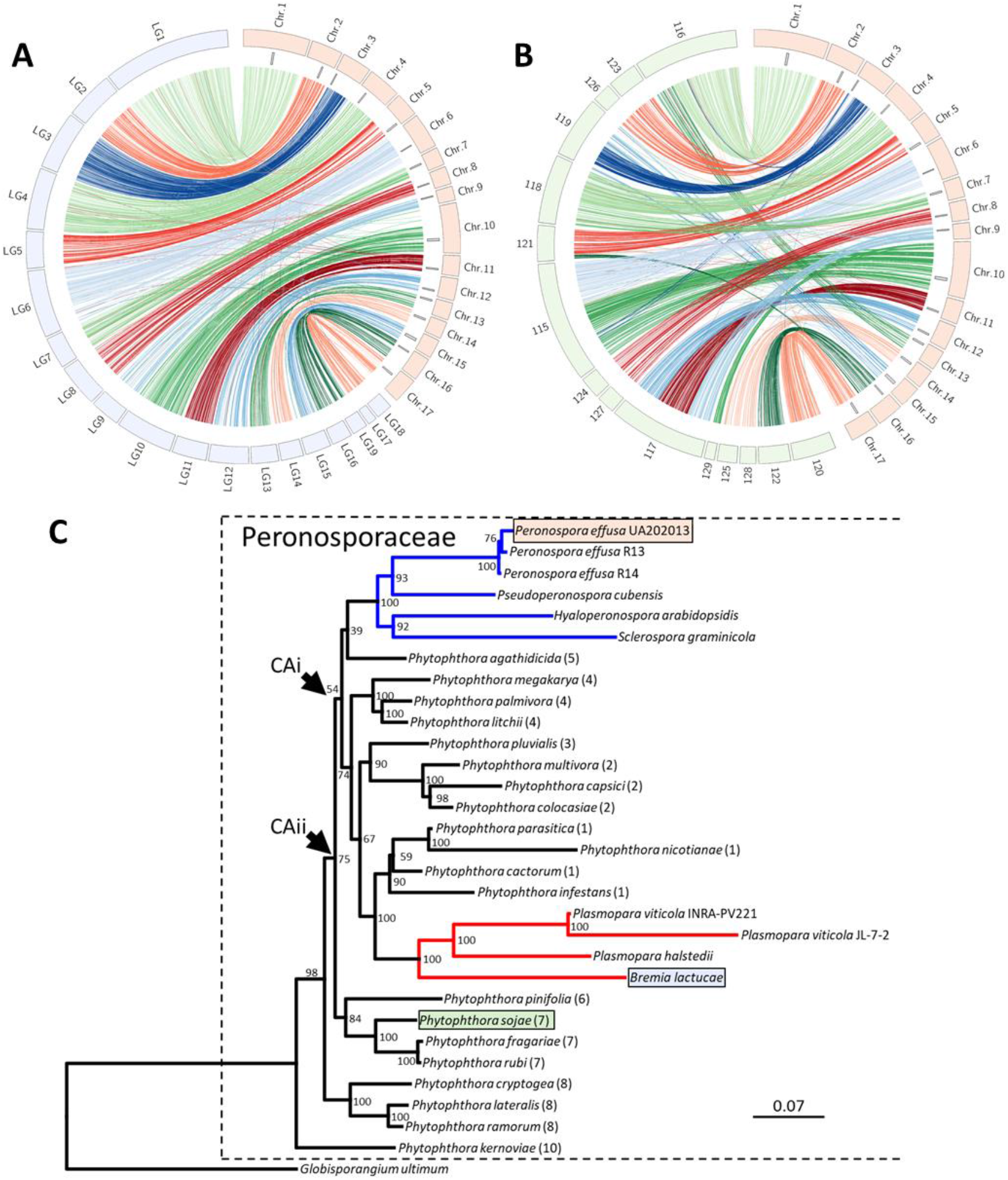
The high synteny of *Peronospora effusa* chromosomes with other oomycetes. Synteny is demonstrated by linking orthologous genes, coloured by *P. effusa* (red) chromosome assignment against A) nineteen genetically oriented scaffolds of the downy mildew *B. lactucae* (blue) and B) fifteen scaffolds larger than 1 Mb of the hemi-biotroph *P. sojae* (green). The plots are scaled to reflect the physical sizes of the chromosomal sequences. C) Maximum likelihood phylogenetic tree calculated from the concatenated alignment of 18 BUSCO proteins surveyed in 31 assemblies of 27 Peronosporaceae spp. with *Globisporangium ultimum* (Pythiaceae) as an outgroup; multiple assemblies for *P. effusa* and *P. viticola* were used. Branches coloured red and blue indicate the two polyphyletic downy mildew clades. Species for which synteny was plotted are highlighted using the same colour palette as in A and B. Parenthesized numbers indicate the *Phytophthora* clade a species has been assigned to. The most recent common ancestor of *P. effusa* and *B. lactucae* is inferred as common to all downy mildews and *Phytophthora* clades 1 to 5 (arrowed CAi). The most recent common ancestor of *P. effusa* and *P. sojae* is inferred to be deeper in the phylogeny (arrowed CAii). Scale indicates the mean number of amino acid substitutions per site. Branch support was calculated from 1,000 bootstraps. This figure was updated from Fletcher *et al*. 2019, Figure 3.

## Discussion

This report describes the first T2T assembly of an oomycete. Sixteen of the 17 chromosomes of *P. effusa* are gapless; only Chr. 15 contains one gap, which likely encodes a highly repetitive rDNA array. This assembly will be a key resource for comparative genomics with all other oomycetes and will advance the study of many important pathogens.

Comparative genomics revealed a high level of synteny between *P. effusa* and the genetically orientated genome assembly of *B. lactucae* (Fletcher et al., 2020). The *P. effusa* chromosomes were numbered to match the corresponding chromosome-scale scaffolds of *B. lactucae*, reported previously (Fletcher et al., 2020), because they have highly conserved single-copy gene contents, similar gene orders along their chromosomes, and likely the same number of chromosomes (Fig. 4A). Therefore, the common ancestor of *P. effusa* and *B. lactucae* may have had a similar chromosomal architecture. Phylogenetic analysis indicated that downy mildews are polyphyletic (Fig. 4C); the most recent common ancestor of biotrophic *P. effusa* and *B. lactucae* is also common to every downy mildew, as well as *Phytophthora* clades 1 through 5 (Bourret et al., 2018; Fletcher et al., 2018; Fletcher et al., 2019). Given the high levels of synteny with *P. sojae*, which is a clade 7 species, the proposed ancestral architecture for many chromosomes is likely to be rooted deeper in the oomycete phylogeny than the last common ancestor of downy mildews (Fig. 4C). Consequently, many downy mildew and *Phytophthora* species may have the same ancestral chromosome configurations; therefore, this T2T assembly of *P. effusa* will serve as a foundational reference for the study of Peronosporaceae and other oomycetes.

Structural variation can be seen around the centromeres when comparing *P. effusa* to *P. sojae* (Supplementary Fig. 1) and in the sub-telomeric region of Chr. 2 between *P. effusa, P. viticola*, and *P. sojae* (Fig. 2). The major structural variation observed between *P. effusa* and *P. sojae* (Fig. 4B) may be due to misassemblies in *P. sojae* or unique structural rearrangements in the evolutionary history of *P. sojae*. Interestingly, one misassembly in the *P. sojae* assembly that joins sequences syntenic to *P. effusa* Chr. 6 and Chr. 10 has previously been reported (Fang et al., 2020). The generation of additional chromosome-scale assemblies will be fundamental in determining how much structural variation exists within and between oomycete species and this T2T assembly of *P. effusa* provides the foundation for such investigations.

This new genome assembly of *P. effusa* was used to analyse the accuracy of previous *P. effusa* draft assemblies. A striking difference is the size of the assemblies, with isolate UA202013 being nearly double that of previous assemblies (Table 3). The increase in assembly size is attributed to the inability of short-reads and high-error rate of long reads to assemble high identity repeats and paralogs (Supplementary Fig. 12, 13). High identity repeats were collapsed, misassembled, or absent in previous short-read assemblies. Collapsed repeat sequences are reported to be present in genome assemblies of other oomycetes (Fletcher et al., 2019) and may result in unreliable analysis, including analysing bursts of LTR activity by nucleotide variation. Calculated LTR divergence since insertion on the assembly of *P. effusa* UA202013 compared to R13 and R14 (generated from short-reads) showed that the resulting profiles differed between long- and short-read assemblies (Supplementary Fig. 4). This result is consistent with short-read assemblies generating chimeric LTR-RTs with inflated divergence in their LTRs. Therefore, estimations of LTR divergence since insertion in oomycete assemblies generated from short-reads should be treated with care (Fletcher et al., 2019).

Short-read assemblies of downy mildews are valuable in identifying most but not all of the gene space. The number of annotated genes for *P. effusa* isolate UA202013 is only slightly higher than isolates R13 and R14 (9,745, 8,607, and 8,571 respectively). More genes were annotated for isolate Pfs1 (Table 3), although many genes were reported to overlap repeats for Pfs1 (Klein et al., 2020); our identification of conserved domains characteristic of retrotransposons in the Pfs1 annotations confirmed the inclusion of LTR-RTs in gene annotations. Orthology analysis demonstrated that the increased gene count for UA202013 was due to improved resolution of paralogs (Supplementary Fig. 12, 13) and not due to the absence of single-copy genes from previous assemblies. These previously collapsed paralogs included putative crinkler proteins, explaining the increase of this class of effector. Therefore, the generation of short-read oomycete assemblies is still informative for comparative analyses because most single-copy genes are recovered but paralogs may be underrepresented.

Short reads are also a valuable resource for comparative analysis between isolates to identify candidate regions containing molecular determinants of race phenotypes. Aligning reads of five other isolates showed that read depth, nucleotide variation, and heterozygosity correlated genome-wide with race phenotypes (Supplementary Fig. 14, 17, and 18). The two race 13 isolates sequenced independently across two studies are very similar, as are the two race 14 isolates. The SNP compositions of these isolates are consistent with somatic derivation from single founder genotypes. Race 14 was proposed to be derived from race 12 after a LOH event (Lyon et al., 2016). Our data support this and favour a LOH event in a region of Chr. 6, resulting in the loss of avirulence alleles (Supplementary Fig. 19). Interestingly, the two race 14 isolates have different lengths of homozygosity, suggesting that they were independent LOH events. Independent LOH events are consistent with Pfs12 (race 12 rather than race 14) and Pfs14 clustering closer to one another than Pfs14 to R14 (Supplementary Fig. 17).

Read-depth analysis also highlighted a cluster of high-identity LWY effectors of Chr. 14 that are unique to UA202013. It is possible that the molecular determinant for UA202013 is located in this cluster and is absent in races 12, 13, and 14. Therefore, these effectors may be recognized by spinach cultivars Clermont, Campania, or Lazio, which all carry the *RPF4* locus. Functional studies are required to test this hypothesis further. Other regions of the UA202013 assembly also had low read depth in the other isolates but did not overlap clusters of effectors (Supplementary Fig. 14).

The T2T assembly demonstrated that some high identity effectors were arranged as tight clusters in the genome (Fig. 1, Supplementary Fig. 5). However, RXLR-EER-WY proteins clustered on the same chromosome do not share 100% identity (Supplementary Fig. 6). Clusters of RXLR and CRN effectors have been described on unplaced scaffolds of *Phytophthora* spp. (Schornack et al., 2009) Therefore, clusters of effectors are likely in other Peronosporaceae species, although they had not been reported in the previous fragmented assemblies of *Peronospora* spp. Such clusters may have consequences for the evolution of new virulence phenotypes because recombination within tightly linked clusters, such as crinkler clusters, will be less frequent than for unlinked genes. The larger clusters of RXLR-EER-WY effectors on the smaller chromosomes may have a frequency of recombination because there will be a higher density of crossovers on the smaller chromosomes. More frequent recombination within and between genes encoding effectors may result in the rapid evolution of novel virulence phenotypes.

In contrast to clusters of genes encoding RXLR-EER-WY effectors, CRNs were tightly clustered on larger chromosomes. CRNs within a cluster were more similar than clustered RXLR-EER-WY effectors (Supplementary Fig. 6). Lower divergence implies that paralogous CRN encoding genes are diversifying more slowly than RXLR-EER-WY encoding genes (Dong and Ma, 2021), reflecting different selection pressures. Interestingly, transient expression of 15 *P. sojae* CRNs in *Nicotiana benthamiana* showed that relatively few of the tested crinklers induced plant cell death (PCD); instead, most crinklers suppressed PCD, consistent with CRNs acting as PCD regulators, not inducers (Shen et al., 2013; Amaro et al., 2017).

The T2T genome assembly allowed an efficient search for horizontal gene transfer (HGT) events by identifying an insertion of genes that are unique to *P. effusa* that lacked orthology with annotations of other oomycete species. Several instances of HGT and subsequent duplication have been previously described for the oomycetes, identifying HGT from fungi or bacteria to oomycetes (Belbahri et al., 2008; Savory et al., 2015; McCarthy and Fitzpatrick, 2016). These used comparative analysis of assemblies for multiple oomycete species/isolates and focused on *Phytophthora* and *Pythium* species. Our comparison of a complete genome to assemblies of other oomycetes provided robust evidence for the recent HGT to an ancestor of *P. effusa*. Two genes have been acquired from phytopathogenic fungi on Chr. 2, one of which was duplicated on Chr. 15 (Fig. 2; Supplementary Fig. 7). The genes on Chr. 2 were also found as a pair in several fungal phytopathogens (Fig. 2). These genes were phylogenetically nested within phytopathogenic fungal sequences and were distinct from paralogs that had orthology to other oomycete species (Fig. 2); this provides strong support for HGT from a fungal species to an ancestor of *P. effusa*. The functions of these genes are unknown; the duplicated gene encoded a metallophosphatase domain (MPD). The acquisition and duplication of the MPD encoding gene suggests a selective advantage (Savory et al., 2015). The large inter-genic distances flanking the horizontally transferred genes means they are in the gene-sparse regions of the genome. The homologs ancestral to oomycetes had inter-genic distances less than 500 bp and therefore are in the gene-dense regions of the genome. The two speed-genome hypothesis postulates that genes in gene sparse-regions evolve faster (Dong et al., 2015). Therefore, these horizontally acquired genes may be evolving differently from their homologs with oomycete ancestry.

In summary, the T2T assembly of *P. effusa* provides an important foundation for comparative genomics between oomycete species and for analysis of diversity within *P. effusa*. Future studies will determine how prevalent the 17-chromosome configuration is between diverse members of the Peronosporaceae and whether there are major structural rearrangements in some lineages. It will also be interesting to determine if the size and chromosomal locations of effector clusters vary, especially between downy mildews, whose ancestors may have adjusted the number of effectors in their genome independently while adaptation to biotrophy occurred. Finally, this assembly provides sequences which can be tested through evolutionary, comparative, and functional analyses to determine what role they play in the success of *P. effusa*.

## Methods

### Isolate propagation and virulence phenotyping

Isolate UA202013 of *P. effusa* was collected from the spinach cultivar Sparrow in Thermal, CA on March 23^rd^, 2020, from a commercial production field. The isolate was propagated prior to DNA extraction using a method described previously (Feng et al., 2013). Sporangia were washed off infected leaves with cold filtered water (4°C) using a vortex mixer. The inoculum was diluted to 1×10^5^ spores/ml and 50 ml was fine sprayed on a 2-week-old flat of Viroflay plants, a downy mildew susceptible spinach cultivar. The inoculated plants were put in a growth chamber where all programmable parameters were adjusted to simulate the average spinach-growing environment. The daily temperature slope was programmed to range between 13°C and 18°C and relative humidity was maintained above 90% using manual misting with cold filtered water. Light was on for 12 h per day with 150 μmol/m/s intensity, except for during the first 24 h to induce sporangial germination in the dark. The plants were taken out from the growth chamber after 12 days to harvest sporangia.

Virulence phenotyping of this isolate was performed using differential lines from the International Working Group on Peronospora following a standardized protocol (Feng et al., 2013; Correll and du Toit, 2018; Feng et al., 2018b) with minor modifications. Briefly, 20 to 30 seedlings of each cultivar, two weeks in age, were inoculated. Sporangia were filtered through four layers of cheese cloth, adjusted to a concentration of 10^5^ sporangia/ml, and spray-inoculated on the 11 differential cultivars using a Badger Basic Spray Gun, Model 250. After inoculation, the plants were incubated in a dew chamber for 24 h, followed by a growth chamber for five days, and a final 24 h in the dew chamber to induce sporulation. The cotyledons and true leaves of each plant were evaluated for downy mildew disease symptoms seven days post-inoculation. Disease incidence (%) for each cultivar was assigned based on the number of plants showing chlorosis and sporulation out of the total number evaluated. Susceptible, resistant, and intermediate disease reaction evaluations were given based on disease incidence.

### DNA extraction and library preparation

Spore suspensions were centrifuged at 8,000 *g* for 5 minutes, the supernatant was removed, and pellets frozen at -80°C. DNA was extracted from pellets (approx. 100 mg) using an OmniPrep kit (G-Biosciences, St. Louis, MO) in 1 ml lysis buffer (G-Biosciences) with 10 µl proteinase K and digested for 1 hour at 60°C with periodic mixing to keep spores suspended in the lysis buffer. Library construction and sequencing were done at the UC Davis Genome Center. DNA quantity and quality was verified with the Quantus fluorometer assay (QuantiFluor ONE dsDNA Dye, Promega, Madison, WI, USA) and Femto Pulse® Automated Pulsed Field (Agilent Technologies, Inc, Santa Clara, CA, USA), respectively. Sheared gDNA (700 ng) was prepared with the Low Input DNA SMRTbell Express Template Prep Kit 2.0 (Pacific Biosciences, Menlo Park, CA, USA) and loaded into a single 8M SMRT cell on a Pacific Biosciences Sequel II.

### Genome sequencing and assembly

PacBio HiFi reads were assembled with Hifiasm v0.14 (Cheng et al., 2021), generating 989 contigs, 11 of which were complete chromosomes. A 12^th^ chromosome (Chr. 10) was obtained from a second Hifiasm assembly generating 57 contigs from reads filtered for mapping to oomycete assigned contigs. Preliminary analysis was conducted by transferring annotations from *P. effusa* R13 (Fletcher et al., 2018) onto the new assembly using Liftoff v1.3.0 (Shumate and Salzberg, 2020). Synteny of intermediate assemblies with the genetically oriented assembly of *B. lactucae* (GCA_004359215.2) (Fletcher et al., 2020) was determined using orthologs as calculated by OrthoFinder v2.2.1 (Emms and Kelly, 2015). The initial 12 chromosomes were found to be colinear with *B. lactucae* linkage group-spanning scaffolds. Multiple *P. effusa* contigs were found to be colinear with other *B. lactucae* linkage group-spanning scaffolds. Long reads aligned to these *P. effusa* contigs with minimap2 v2.17-r954-dirty (Li, 2018) were assembled independently again with Hifiasm v0.14 (Cheng et al., 2021) and complete chromosome contigs were assembled for two chromosomes (Chr. 6 and Chr. 11). Chr. 4 and Chr. 16 were closed by aligning contigs generated by Hifiasm with contigs generated by HiCanu v2.0 (Nurk et al., 2020), extending the Hifiasm contigs by 8,619 bp and 12,694 bp, respectively, and adding telomeres to the sequence. Two telomeric contigs of length 2,338,820 bp and 30,638 bp remained in the assembly, both terminating in repetitive rDNA sequences. They could not be aligned, nor could reads be found that bridged the sequences. Read alignment showed that the depth was high, indicating that a large repeat array was present, as such that the gap was bridged with 100 unknown bases (N).

### Genome annotation and comparative analysis

Genome annotation was performed comparably to other *P. effusa* assemblies (Fletcher et al., 2018). A repeat library was defined with RepeatModeler v1.73 (Smit and Hubley, 2008) and masked with RepeatMasker v4.0.9 (Smit et al., 2013). Repeats were annotated in six other *P. effusa* genome assemblies (Feng et al., 2018a; Fletcher et al., 2018; Klein et al., 2020) using the same repeat library. Repeat sequences of UA202013 identified by RepeatMasker were clustered using CD-Hit v4.8.1 (Li and Godzik, 2006), requiring 90% identity to the centroid sequences. Pairwise identity within each cluster was calculated using BLASTn v2.10.1 (Altschul et al., 1990). Inter- and intra-chromosomal clusters were visualized using Circos v0.69-8 (Krzywinski et al., 2009). Nucleotide variation between LTRs assembled in UA202013 was calculated as previously described (Fletcher et al., 2019). Briefly, LTR-RTs were identified and annotated using LTRharvest v1.5.9 (Ellinghaus et al., 2008) and LTRdigest v1.5.9 (Steinbiss et al., 2009), respectively. Internal domains were clustered with VMatch v2.3.0 and within cluster alignments of 5’ and 3’ LTRs were generated with ClustalO v1.2.0 (Sievers et al., 2011). The divergence between LTR pairs was calculated with BaseML v4.7b (Yang, 2007). Previous LTR divergence results for *P. effusa, B. lactucae, P. infestans, P. ramorum*, and *P. sojae* were taken from a previous analysis (Fletcher et al., 2019). Results were plotted using R v4.0.1 base packages (R Development Core Team, 2012).

Genes were annotated with MAKER v2.31.10 (Campbell et al., 2014) using a SNAP v2013-02-16 (Korf, 2004) HMM model generated in a previous study (Fletcher et al., 2018). Additional effectors were predicted from single open reading frames, searching for crinkler, RxLR, and EER motifs and WY domains. Effector annotations were integrated with the MAKER annotations to ensure no overlapping gene models were introduced. Annotations were validated through orthology analysis with annotations of 37 other oomycetes using OrthoFinder v2.2.1 (Emms and Kelly, 2015). Intergenic distances were calculated from the GFF file.

Effector gene clusters were identified by clustering effector amino acid sequences using CD-Hit v4.8.1 (Li and Godzik, 2006), requiring 40% identity to the centroid sequences for RXLR-EER/WY proteins, 70% for crinklers. Pairwise alignments within each cluster were calculated with BLASTp v2.10.1 (Altschul et al., 1990). Links were visualized with Circos v0.69-8 (Krzywinski et al., 2009). An effector gene cluster was manually called if the cluster contained three or more annotations on the same chromosome. Overlapping clusters were merged into a single cluster.

Expanded paralogs in UA202013 were identified by contrasting orthogroup gene counts, calculated with OrthoFinder v2.2.1 (Emms and Kelly, 2015), for UA202013 with R13 and R14. These results were taken from the larger analysis containing annotations from 38 genome assemblies. Orthogroups in which UA202013 contained five more genes than either R13 or R14 were considered expanded. Read depth of the annotations in these orthogroups was calculated by aligning reads back to their respective assemblies using bwa v0.7.17 (Li, 2013) or minimap2 v2.17-r954-dirty (Li, 2018) and calculated with SAMtools v1.11 bedcov (Li et al., 2009). Stacked bar charts and dotplots were generated using R v4.0.1 (R Development Core Team, 2012) and ggplot2 v3.3.3 (Wickham, 2016). Completeness was calculate using BUSCO v2.0 with the protists_odb9 dataset (Simao et al., 2015). Translated single copy orthologs were added to the 18-BUSCO-gene concatenated alignment previously reported (Fletcher et al., 2019). Protein alignments for each gene were re-calculated independent of one another using MAFFT v7.245 (Katoh and Standley, 2013). Alignments were concatenated and a single tree produced using RAxML v8.2.9 (Stamatakis, 2014), with 1,000 bootstraps and the PROTGAMMAAUTO model. The tree was visualized in Geneious v8.0.5 (Kearse et al., 2012).

Horizontal gene transfer was investigated in the genes unique to *P. effusa* isolate UA202013 using BLASTp v2.10.1 (Altschul et al., 1990) to search the NCBI nr database excluding oomycetes. Genes with a low e-value were tested phylogenetically. A ClustalW alignment was built containing the candidate HGT gene and top BLASTp hits from non-oomycete and oomycete species. This alignment was used to build a neighbour-joining consensus tree to determine if the candidate gene was nested in non-oomycete sequences. A phylogenetic analysis was performed and visualized using Geneious v8.0.5 (Kearse et al., 2012). The same analysis was performed on flanking genes. Orthology of flanking genes was investigated and plotted for contiguous oomycete genome assemblies using R v4.0.1 (R Development Core Team, 2012), ggplot2 v3.3.3(Wickham, 2016), and ggforce v0.3.3 (Pedersen, 2019). Chromosomal fragments of *P. effusa* Chr. 2 and Chr. 15 containing a duplicated gene were aligned against each other using BLASTn. The alignment results were visualized using R v4.0.1 (R Development Core Team, 2012) and ggplot2 v3.3.3 (Wickham, 2016).

Synteny was plotted for the final *P. effusa* against *B. lactucae* and *P. sojae* by linking single copy orthologs between the assemblies with Circos v0.69-8 (Krzywinski et al., 2009). Twelve putative *P. effusa* centromeres were inferred through synteny with the more fragmented *P. sojae* assembly GCA_009848525.1 for which coordinates of 12 centromeres have been previously defined (Fang et al., 2020). An additional five putative centromeres (Chr. 5, Chr. 7, Chr. 9, Chr. 12, and Chr. 13) were defined by identifying BLASTn hits (alignment length >500 bp) of a *P. sojae* CoLT previously hypothesized as diagnostic of oomycete centromeres (Fang et al., 2020). This alignment criterion was relaxed in comparison to similar surveys searching for *B. lactucae* and *P. citiricola* centromeres (Fang et al., 2020).

Previous *P. effusa* genome assemblies (Feng et al., 2018a; Fletcher et al., 2020; Klein et al., 2020) were aligned to the new genome assembly using BLASTn (Altschul et al., 1990), filtering for alignments with a minimum length of 2,500 bp and scoring ≥95% identity. Genes for which 95% of the feature was within assembly-to-assembly alignments were reported as covered, regardless of the alignment length. Long reads of isolate UA202013 (this study) were aligned back to the assembly with Minimap2 v2.17-r954-dirty (Li, 2018). Short reads of R13, R14 (Fletcher et al., 2018), pfs12, pfs13, and pfs14 (Feng et al., 2018a) were aligned with bwa v0.7.17 mem (Li, 2013). Variants for long and short read alignments were called with FreeBayes v1.3.1-17-gaa2ace8 (Garrison and Marth, 2012). Bi-allelic single nucleotide polymorphisms (SNPs) were used to cluster isolates genome-wide and per chromosome. Homozygous references were coded as 0, heterozygous as 0.5, and homozygous alternative as 1. Euclidean distances between isolates were calculated using R v4.0.1 base function dist()(R Development Core Team, 2012). Results were visualized using Heatmap2 from the R package gplots v3.1.1 (Warnes et al., 2020).

For visualization, the genome was broken into 100 kb windows with a 25 kb step. The AT of each window was calculated using BEDTools2 v2.29.2 nuc (Quinlan, 2014). The density of repeats, genes, structural variants (UA202013 only), and SNPs was calculated using BEDTools2 v2.29.2 (Quinlan, 2014). The normalized read depth and coverage of previous draft genome assemblies per window were calculated using BEDTools2 v2.29.2 (Quinlan, 2014). Per window heterozygosity was calculated as the number of SNPs genotyped 0/1 by FreeBayes v1.3.1-17-gaa2ace8. All tracks were formatted as white-space separated files and plotted using Circos v0.69-8 (Krzywinski et al., 2009).

## Acknowledgements

We thank A. Garcia-Llanos (UC Davis) and A. Achieta (USDA/ARS, Salinas) for DNA extractions, O. Nguyen (UC Davis) for library preparation and sequencing, H. Xu (UC Davis) for raw data submissions to NCBI, and E. Georgian (UC Davis) for editorial services. The sequencing was carried out by the DNA Technologies and Expression Analysis Cores at the UC Davis Genome Center, supported by NIH Shared Instrumentation Grant 1S10OD010786-01. The bioinformatic analysis was carried out using the UC Davis LSSC0 High Performance Computing cluster maintained by the UC Davis Bioinformatics Core.

## Author Contributions

KF performed the bioinformatic analysis and drafted the manuscript. AP collected the isolate. OS, JC, CF, SK, and KC conducted the phenotyping of the isolate. AV conceived, coordinated, and provided funding for sequencing. RM contributed to the bioinformatic analyses and to all drafts. All authors contributed to the final manuscript and approved the submission.

## Data availability

The raw reads, genome assembly, and annotation are available under NCBI BioProject PRJNA745455. All software used is described and cited in the Methods.

