## Supplementary Figures 1 to 20 for "Ancestral chromosomes for the Peronosporaceae inferred from a telomere-to-telomere genome assembly of *Peronospora effusa*"

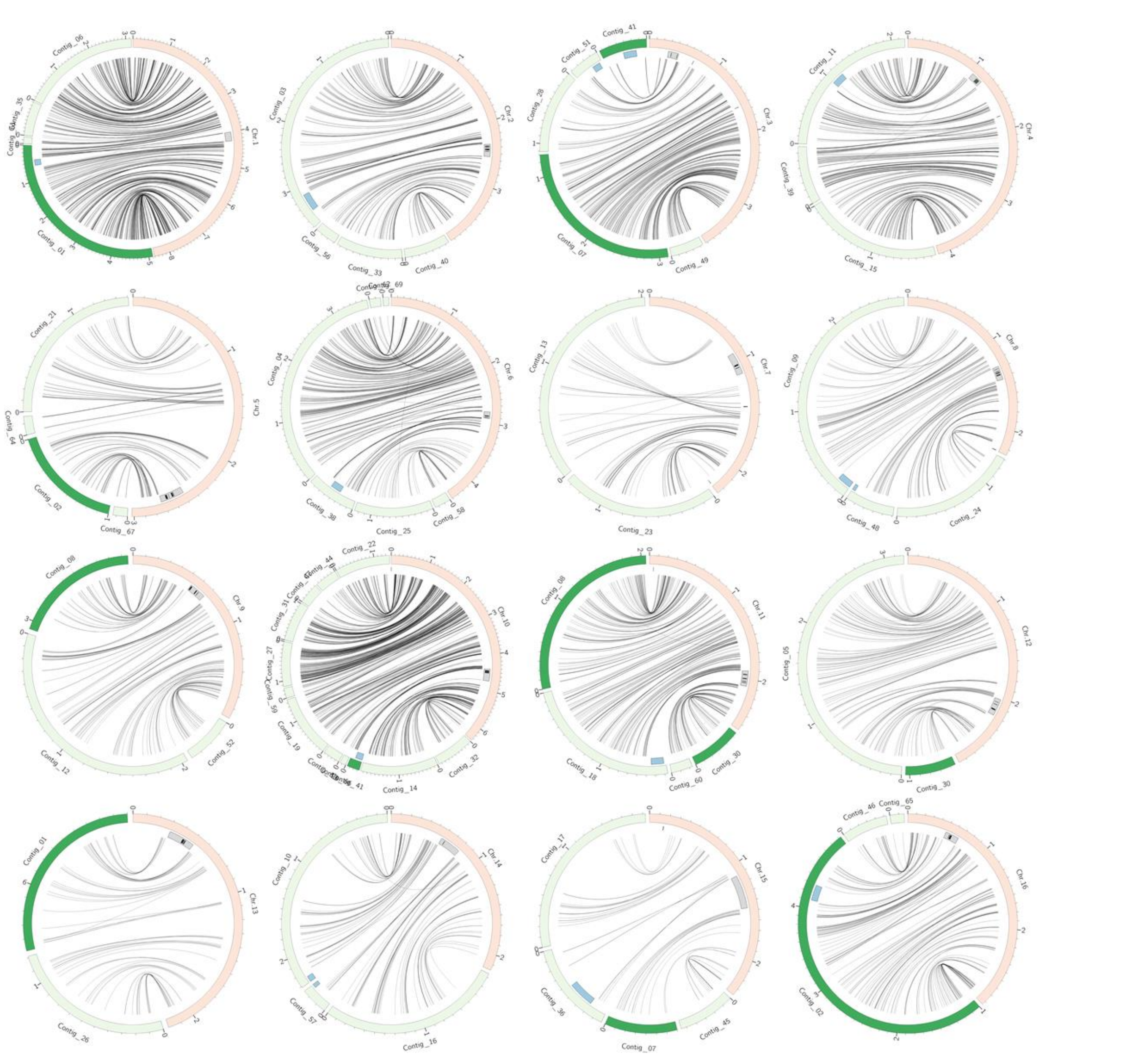



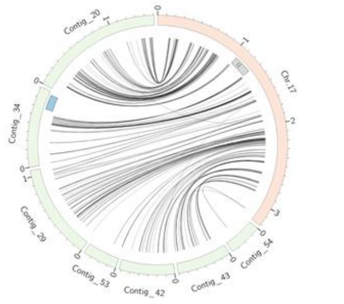



**Supplementary Fig. 1. Orthology of every *Peronospora effusa* chromosome (pink) with contigs of *P. sojae* v2019.1 (GCA_009848525.1; green).** Dark green contigs indicate contigs of *P. sojae* v2019 that align to multiple chromosomes of *P. effusa*. Centromere coordinates previously reported for *P. sojae* are shown in blue, putative *P. effusa* centromere coordinates in grey. Black marks indicated the presence of a Copia-like transposon (CoLT) element in *P. effusa*. CoLT elements have previously been reported as enriched in the centromeres of *P. sojae*, *P. citricola*, and *B. lactucae*; at least one CoLT element was detected in the putative *P. effusa* centromeres of all chromosomes, except Chr. 1 and Chr. 15. Twelve putative centromeres of *P. effusa* were syntenic with centromeres of *P. sojae*, though some gene rearrangements were inferred across and adjacent to centromeres (Chr. 1, Chr. 6, Chr. 7, Chr. 8, and Chr. 15). Five putative centromeres of *P. effusa* were defined by the presence of CoLT elements (Chr. 5, Chr. 7, Chr. 9, Chr. 12, and Chr. 13).

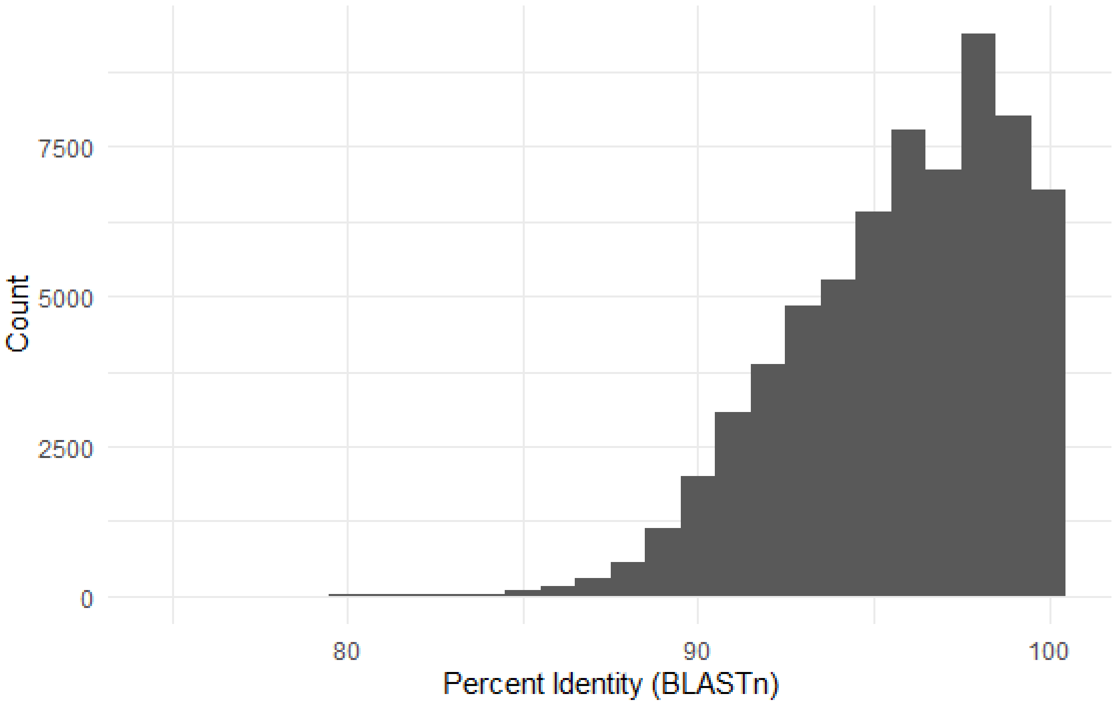


**Supplementary Fig. 2. Pairwise identity of clustered repeat sequences in *P. effusa*.** Sequences annotated as repeats were clustered using CD-hit (90% to centroid sequence). Sequences assigned to each cluster were aligned to one another using BLASTn. The percent identities for 66,858 pairwise alignments were plotted as a histogram. This demonstrates that the genome of *P. effusa* contains thousands of repetitive elements with ≥98% identity between them, which is difficult to resolve with short read sequencing.

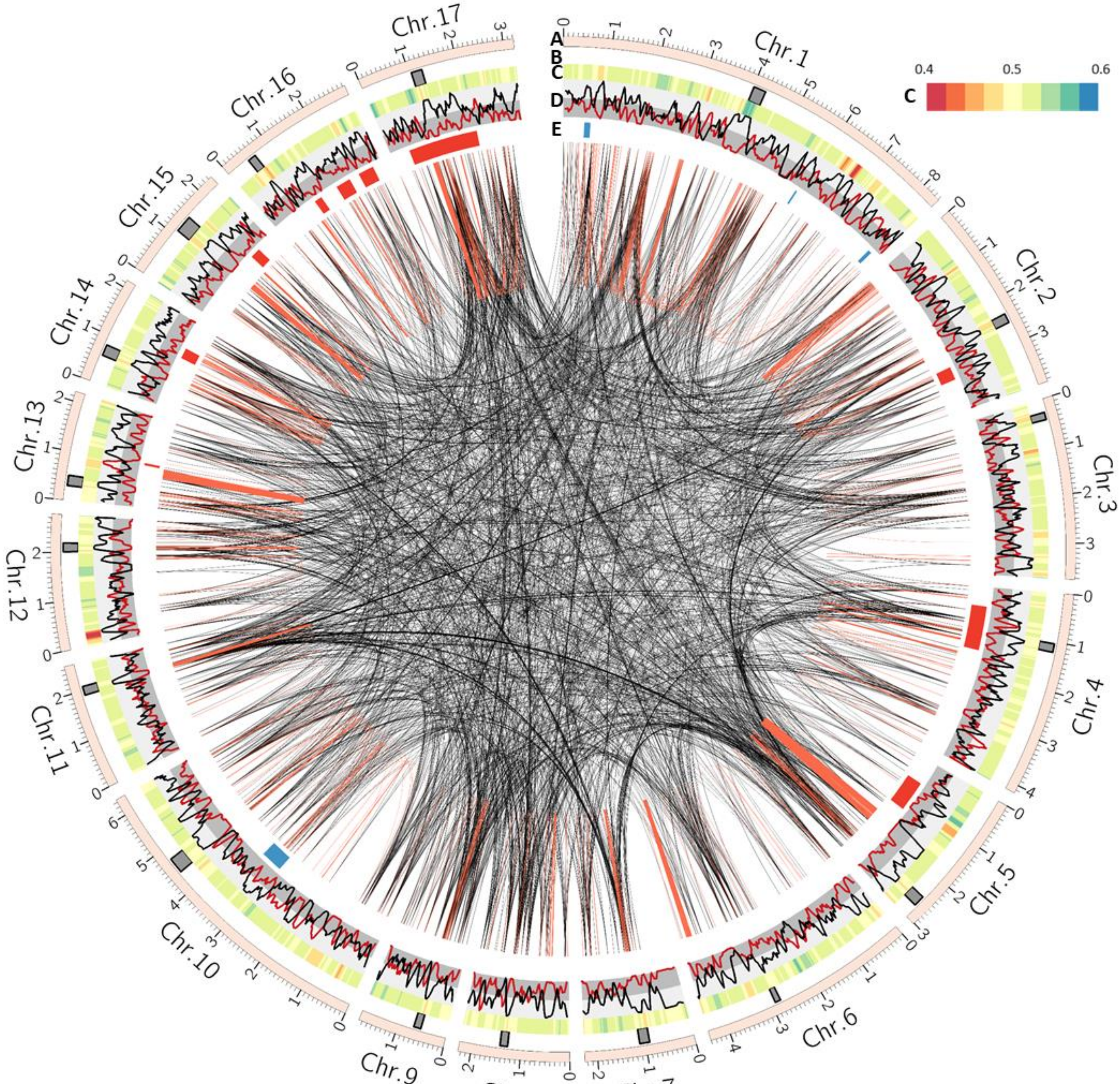


**Supplementary Fig. 3. Genomic distribution of high identity repetitive sequences *of P. effusa*.** Links were drawn between repeats with a pairwise identity greater than 97.5%. In total, 24,159 links are plotted, 21,781 are intra-chromosomal (red), and 2,378 are inter-chromosomal (black). High identity repeats are widely distributed in the genome. Tracks A–D as Fig. 1. Track E as Fig. 1F.

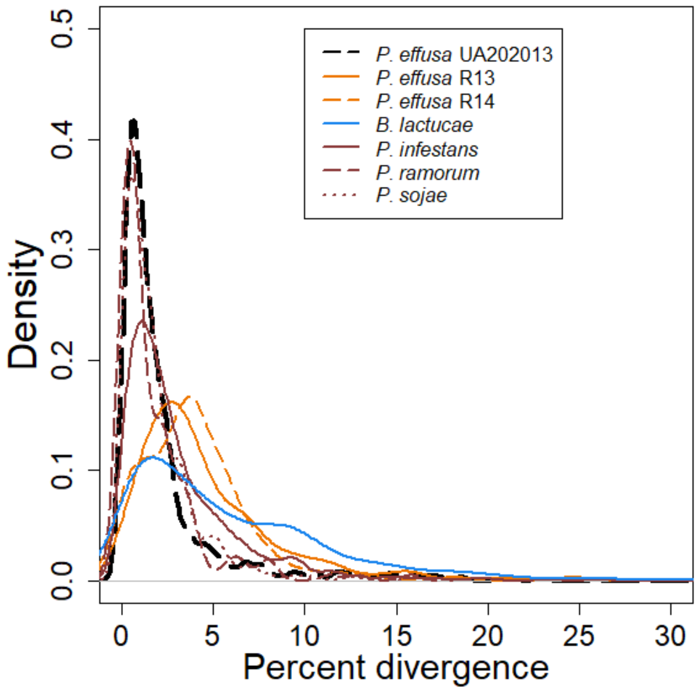


**Supplementary Fig. 4. Estimated age distribution of long-terminal repeat retrotransposons (LTR-RTs) in *P. effusa*.** LTR-RT elements were detected using ltrharvest and annotated using ltrdigest. Full length elements were clustered with vmatch, based on similarity of internal domains. ClustalO was used to align all 5' and 3' LTR sequences within each cluster and divergence between pairs was calculated with BaseML. The density of LTR divergence was plotted in R and compared to results previously reported (Fletcher *et al.* 2019). The assembly of *P. effusa* isolate UA202013 contains a higher density of less diverged LTR elements, possibly be due to chimeric LTR-RTs in previous *P. effusa* assemblies. The density profile of *P. effusa* UA202013 looks more like *P. ramorum* and *P. sojae*, which were assembled with Sanger reads. Therefore, short read assemblies may not be suitable to infer LTR-RT insertion activity in oomycete genome assemblies.

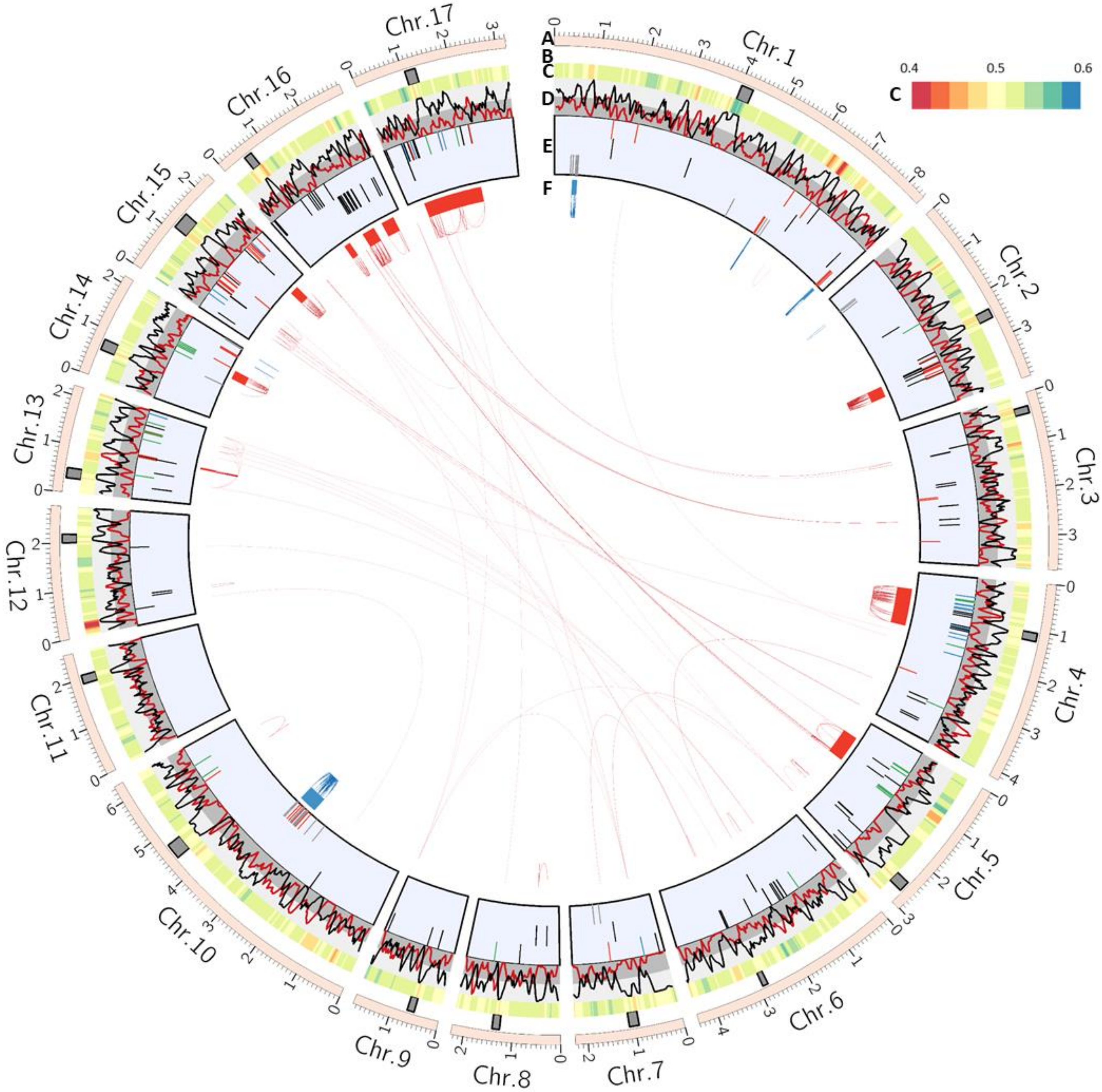


**Supplementary Fig. 5. Organisation of effector proteins into genomic clusters**. Tracks A–F as Fig. 1. The clusters depicted on track F were derived by clustering all RXLR-EER/WY and CRN proteins by identity. Ten clusters of RXLR-EER/WY (red) were identified. These clusters contained three or more RXLR-EER/WY proteins sharing >40% pairwise identity on the same chromosome. This accounted for 99 of 209 RXLR-EER/WY proteins. Four clusters of crinklers (blue) were identified. These clusters contained three or more crinkler proteins sharing >70% pairwise identity. This accounted for 74 of 98 crinkler proteins.

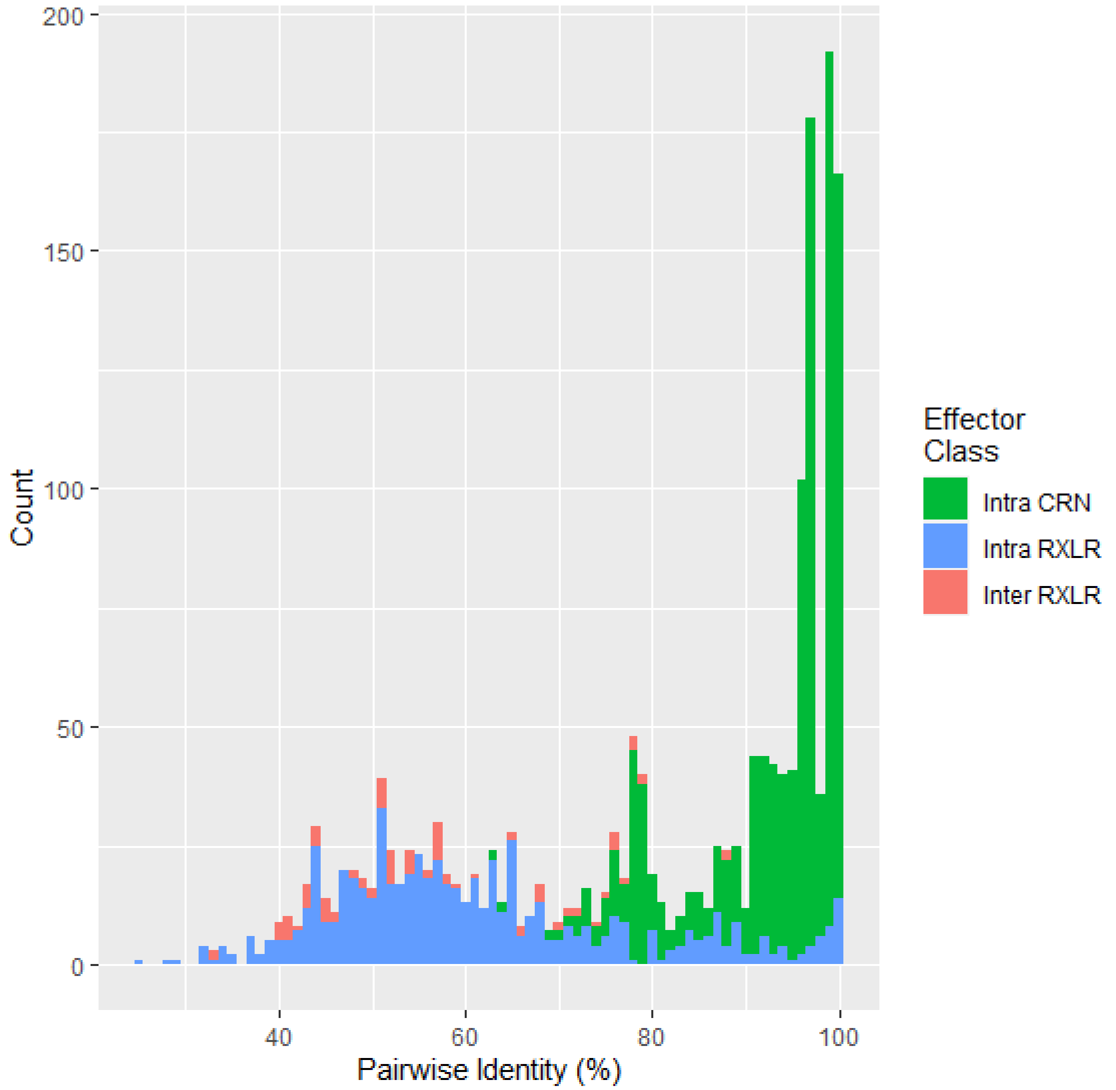


**Supplementary Fig. 6. Pairwise identity calculated by aligning clusters of effector proteins in *P. effusa* with BLASTp, depicted as links in Supplementary Fig. 5.** All crinkler alignments were inter-chromosomal and shared a high percent identity with one another. Most RXLR-EER-WY protein clusters were between intra-chromosomal annotations, although some were inter-chromosomal, both following a similar distribution. High identity (>90%) alignments were only identified between intra-chromosomal comparisons.

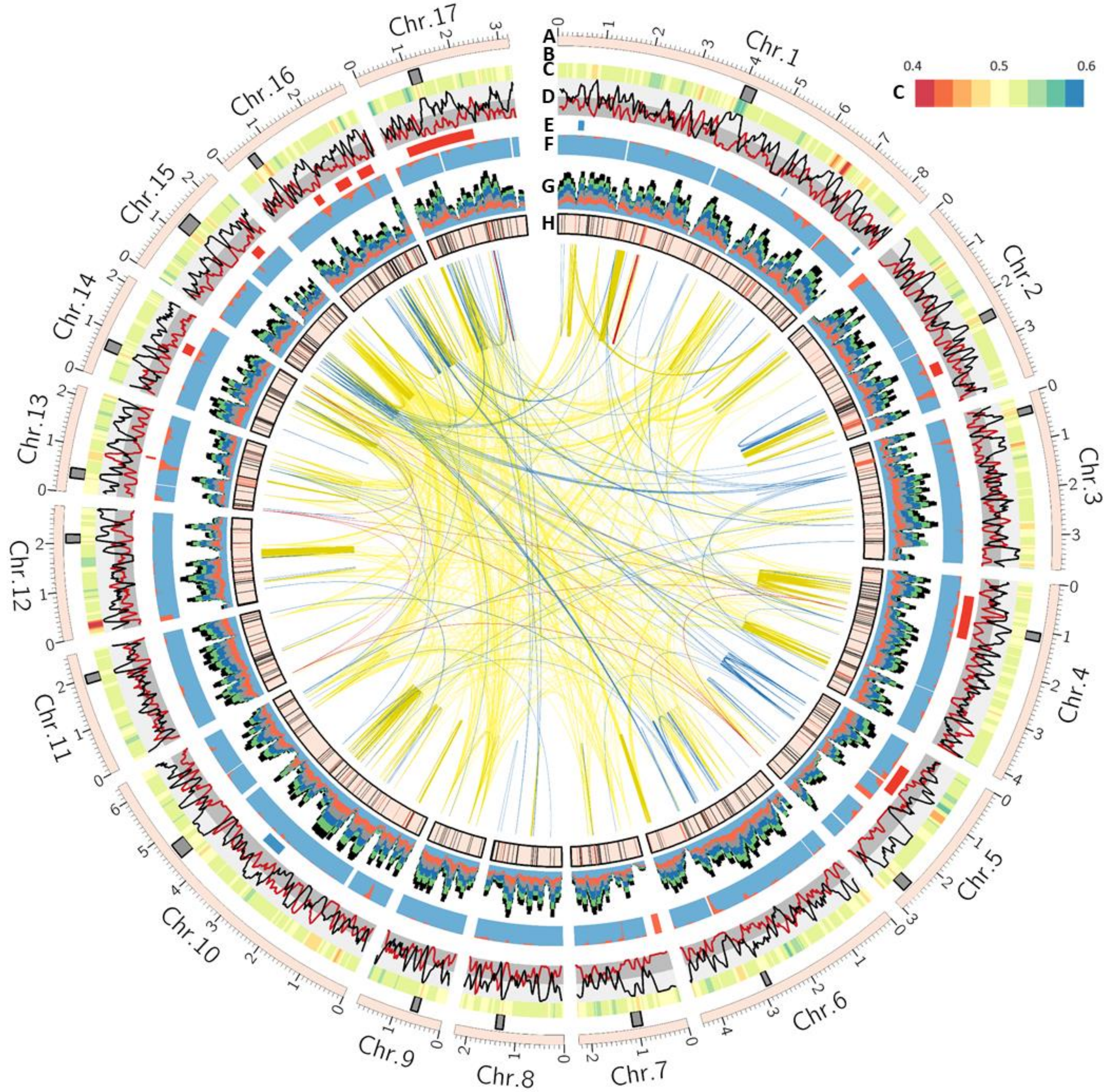


**Supplementary Fig. 7. Orthology between *P. effusa* UA202013 and two sets of annotations from smaller *P. effusa* assemblies.** Tracks A to D are the same a Fig. 1A-D. Track E is Fig. 1F. Track E is Fig. 1F. Track F is Fig. 1G. Track G is Fig. 1H. Track H represents genes unique to *P. effusa*. Black marks indicate *P. effusa* UA202013 annotations also annotated in R13, R14, or both. In total, this included 725 UA202013 annotations. Red marks indicate *P. effusa* UA202013 annotations not annotated in either R13 or R14. This was a total of 281 annotations. Annotations unique to UA202013 can be seen on all chromosomes, except 9. Links indicate paralogs. Yellow links join paralogs for which orthologs could be detected in other oomycetes. Blue links join paralogs for which homologs could be detected in the genome assemblies of *P. effusa* R13 or R14. Red links join paralogs for which homologs were not detected in the genome assemblies of either *P. effusa* R13 or R14.

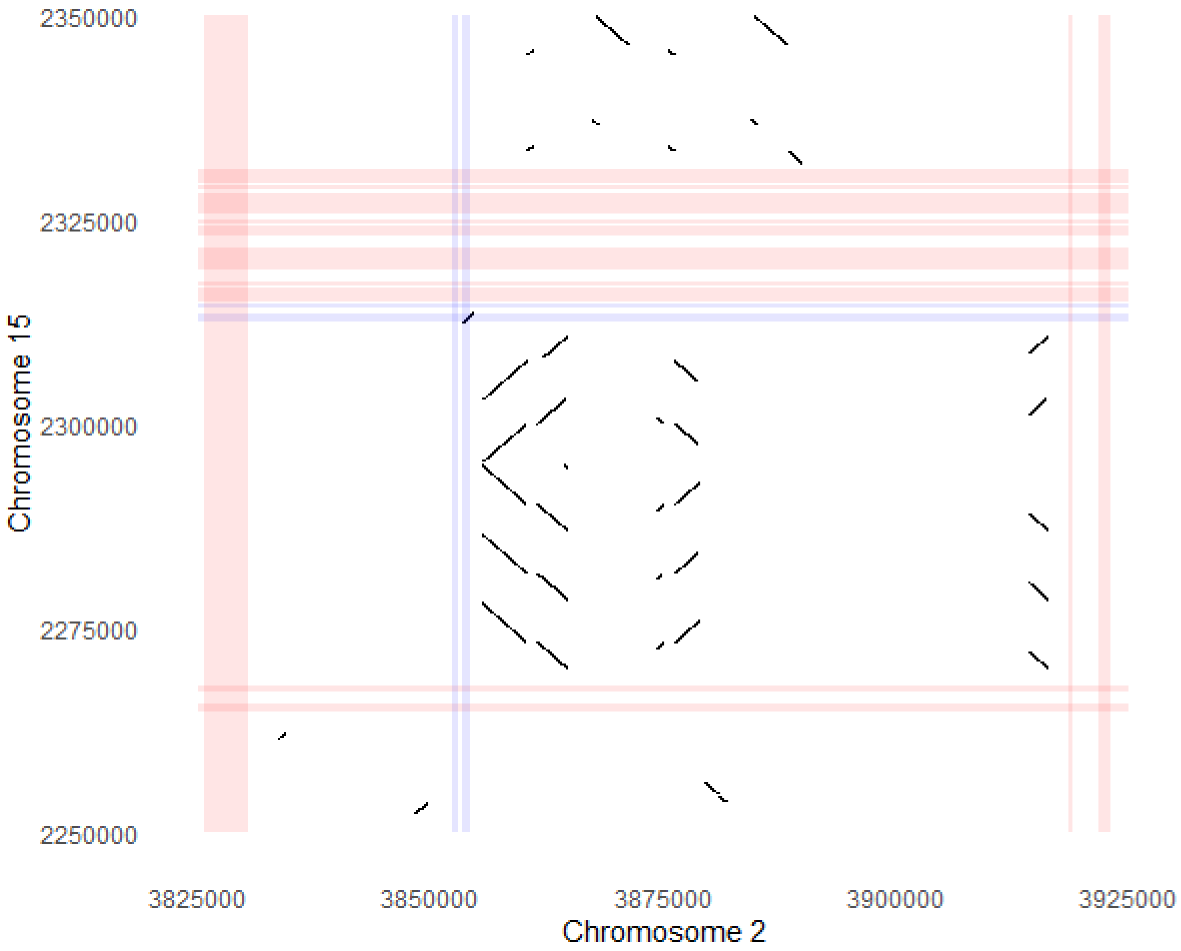


**Supplementary Fig. 8. BLASTn alignment between two regions of *P. effusa* chromosomes containing paralogous genes, likely acquired through horizontal gene transfer (HGT).** Coloured bars indicate genes, red have orthology to other oomycetes, blue have orthology to fungi and may have been acquired by HGT. Black diagonal lines show pairwise alignment between the two regions. On Chr. 2 the HGT acquired gene is in a gene sparse, repeat rich region. The paralog on Chr. 15 is close to a block of oomycete genes and five copies of a repeat sequence present on Chr. 2

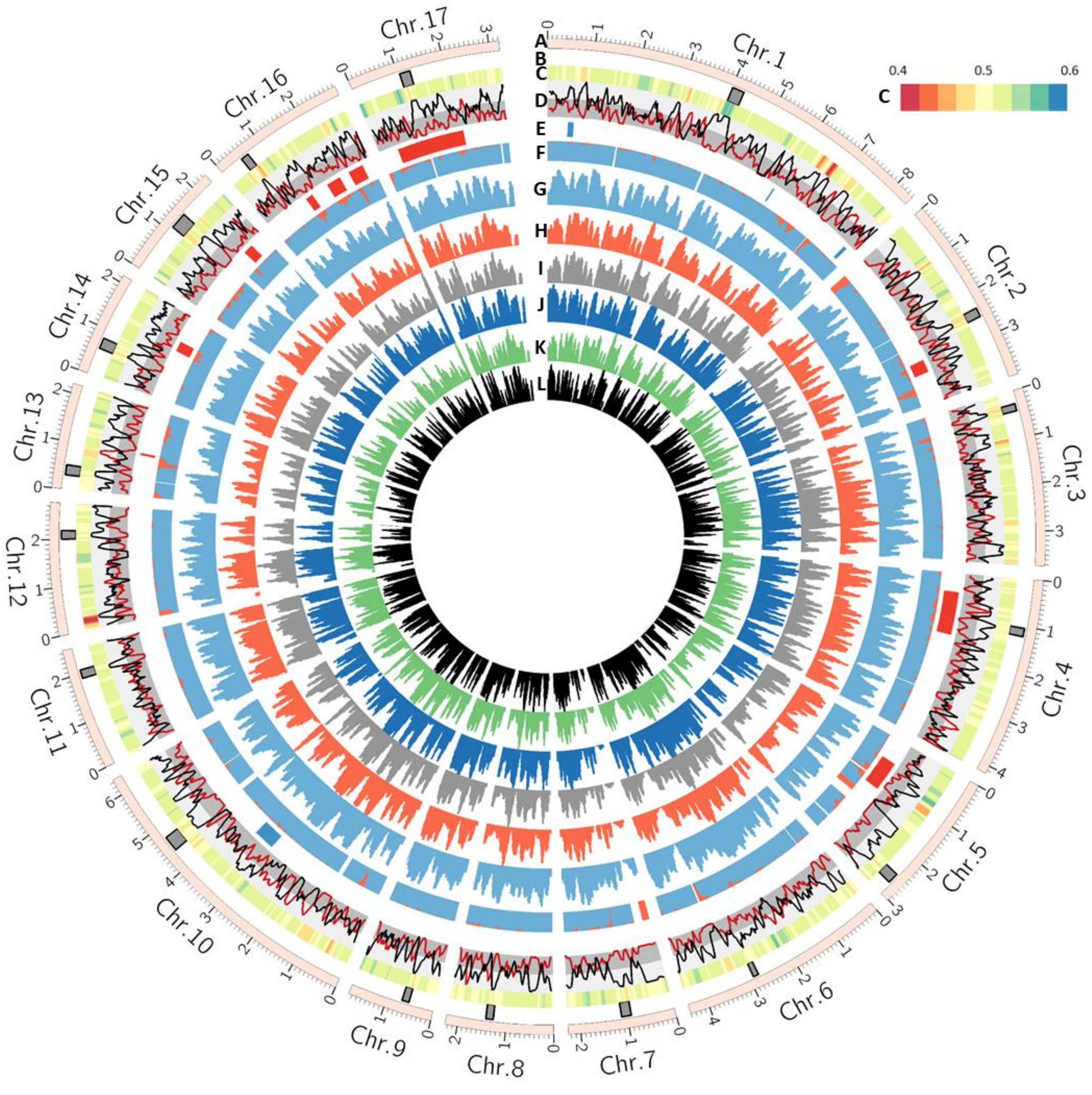


**Supplementary Fig. 9. Coverage of BLASTn alignments between *Peronospora effusa* UA202013 and six smaller *P. effusa* assemblies.** Tracks A to D are the same a Fig. 1A–D. Track E is Fig. 1F. Track F is Fig. 1G. Tracks G to L are an expansion of Fig. 1H. Track G is Pfs1, H is Pfs12, I is Pfs13, J is R13, K is Pfs14, and L is R14. BLASTn alignments were filtered for 95% identity, 2,500 bp minimum alignment length. The number of bases of each *P. effusa* UA202013 100 kb bin covered by alignments to other isolates was calculated and plotted. Multiple and split alignments per contig were considered.

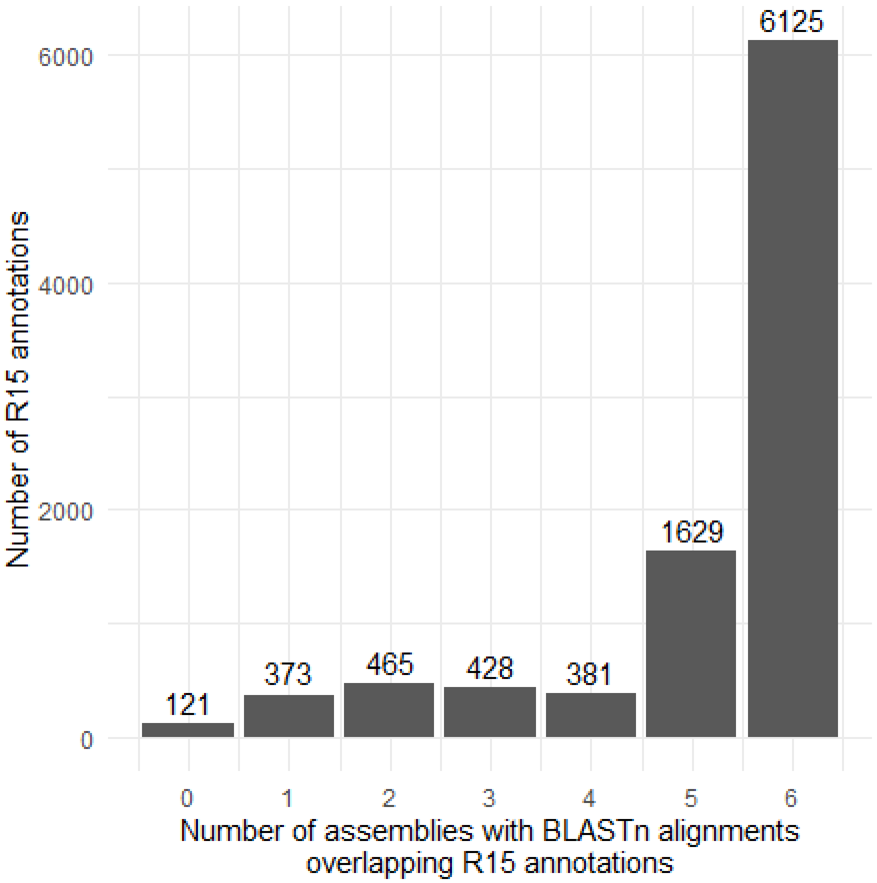


**Supplementary Fig. 10. *P. effusa* UA202013 annotations covered by BLASTn alignments generated with other genome assemblies of *P. effusa*.** To be scored as present in other isolates, the *P. effusa* gene had to be 95% covered by BLASTn alignments. Only 121 *P. effusa* UA202013 annotations were not found in any other *P. effusa* genome, suggesting little novelty. Alignments to the majority of *P. effusa* UA202013 annotations were found in all six other assemblies (Pfs1; Klein *et al.* 2020, Pfs12, Pfs13, Pfs14; Feng *et al.* 2018, R13 and R14; Fletcher *et al.* 2018).

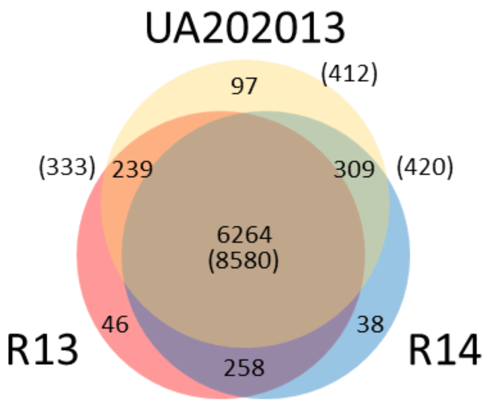


**Supplementary Fig. 11. Three-way Venn diagram illustrating the number of orthogroups common to three *P. effusa* isolates.** Numbers were calculated in a multi-species ortholog analysis and filtered to only consider annotated assemblies of *P. effusa*. Parenthesised numbers indicate the number of UA202013 genes assigned to the orthogroups.

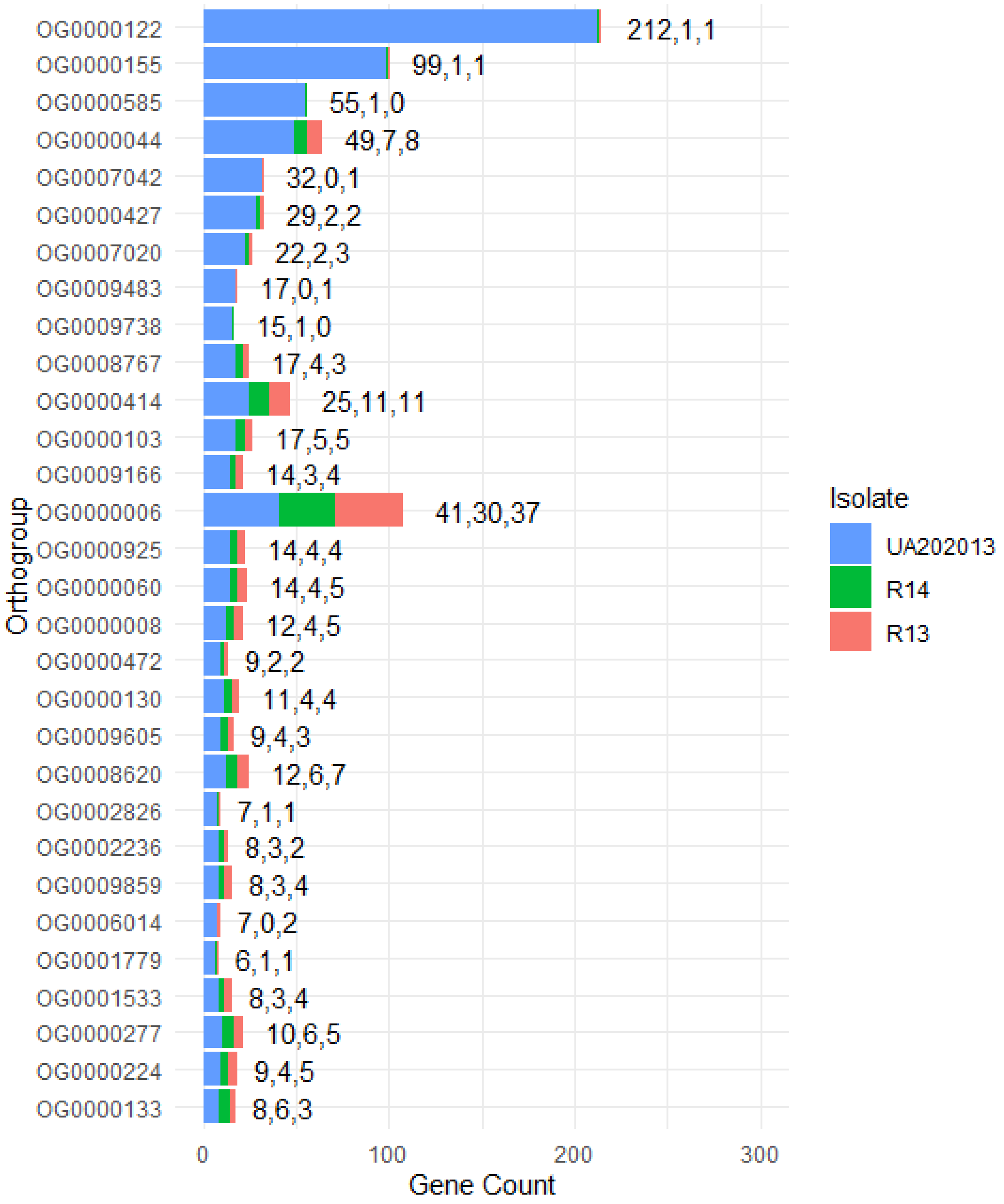


**Supplementary Fig. 12. Assignment of *P. effusa* annotations to orthogroups indicates superior resolution of paralogs in the UA202013 assembly.** The 30 orthogroups for which the largest discrepancy between UA202013 and R13 or R14 are plotted, ordered by the size of discrepancy between the assemblies. Comma separated values indicate the number of genes assigned to the orthogroup for UA202013, R14, and R13. Across all orthogroups, UA202013 had 964 more annotations assigned orthology compared to R13 and 968 compared to R14, suggesting that high identity genes, such as paralogs, are likely collapsed in short read assemblies. In contrast, only four more R14 annotations were assigned orthology than for R13.

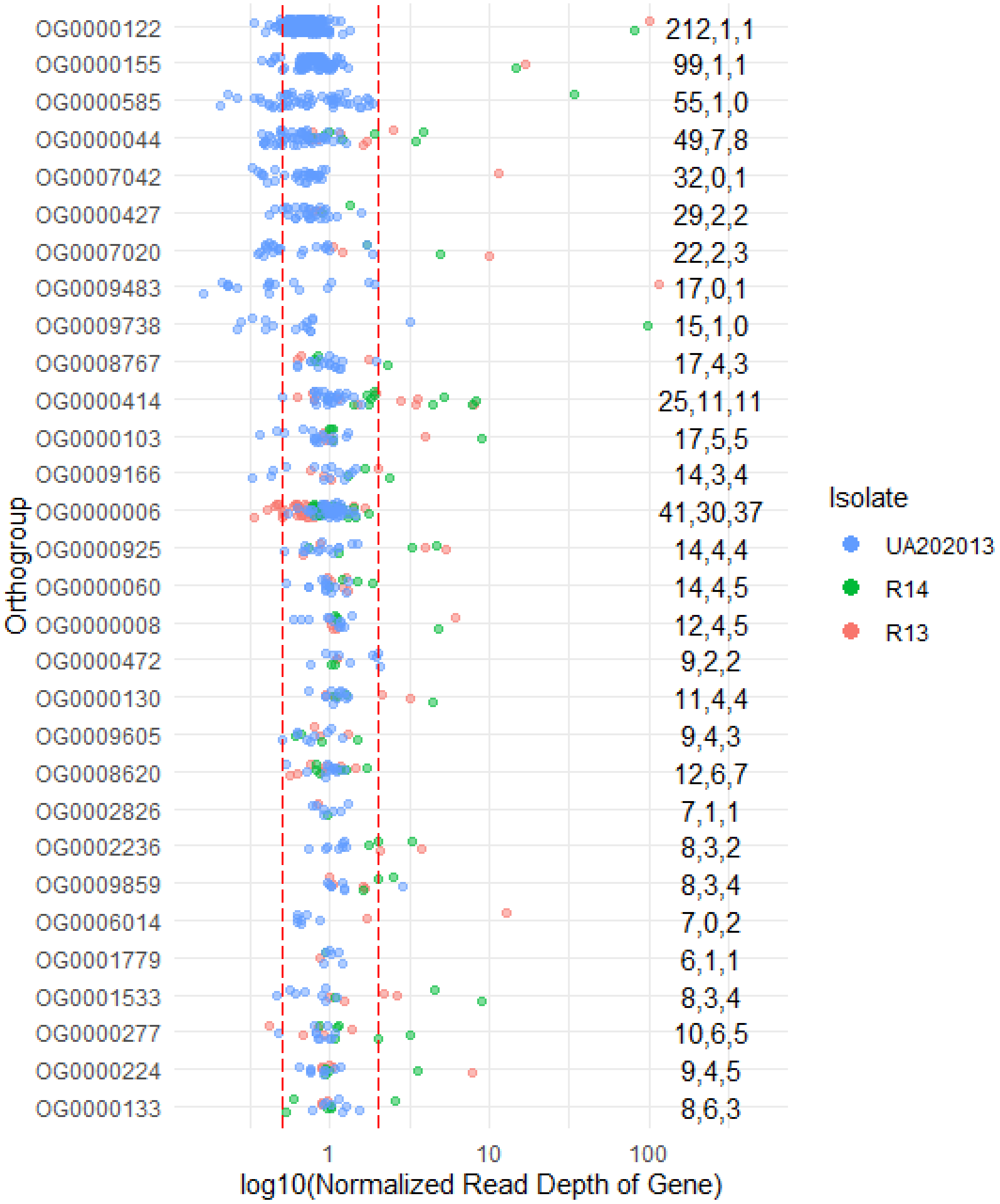


**Supplementary Fig. 13. Read depth of annotations from different *P. effusa* assemblies assigned to common orthogroups identified in Supplementary Fig. 12.** The red dashed lines indicate the boundaries of 0.5x to 2x normalised read depth. Annotations with read depth greater than two are likely to represent high identity paralogs that were collapsed in short read assemblies. For most orthogroups, annotations of R13 and R14 are seen to have higher than expected read depths, indicating that paralogs have been collapsed into a single annotation in the genome assembly. Read depth of UA202013 annotations was typically as expected for every orthogroup.

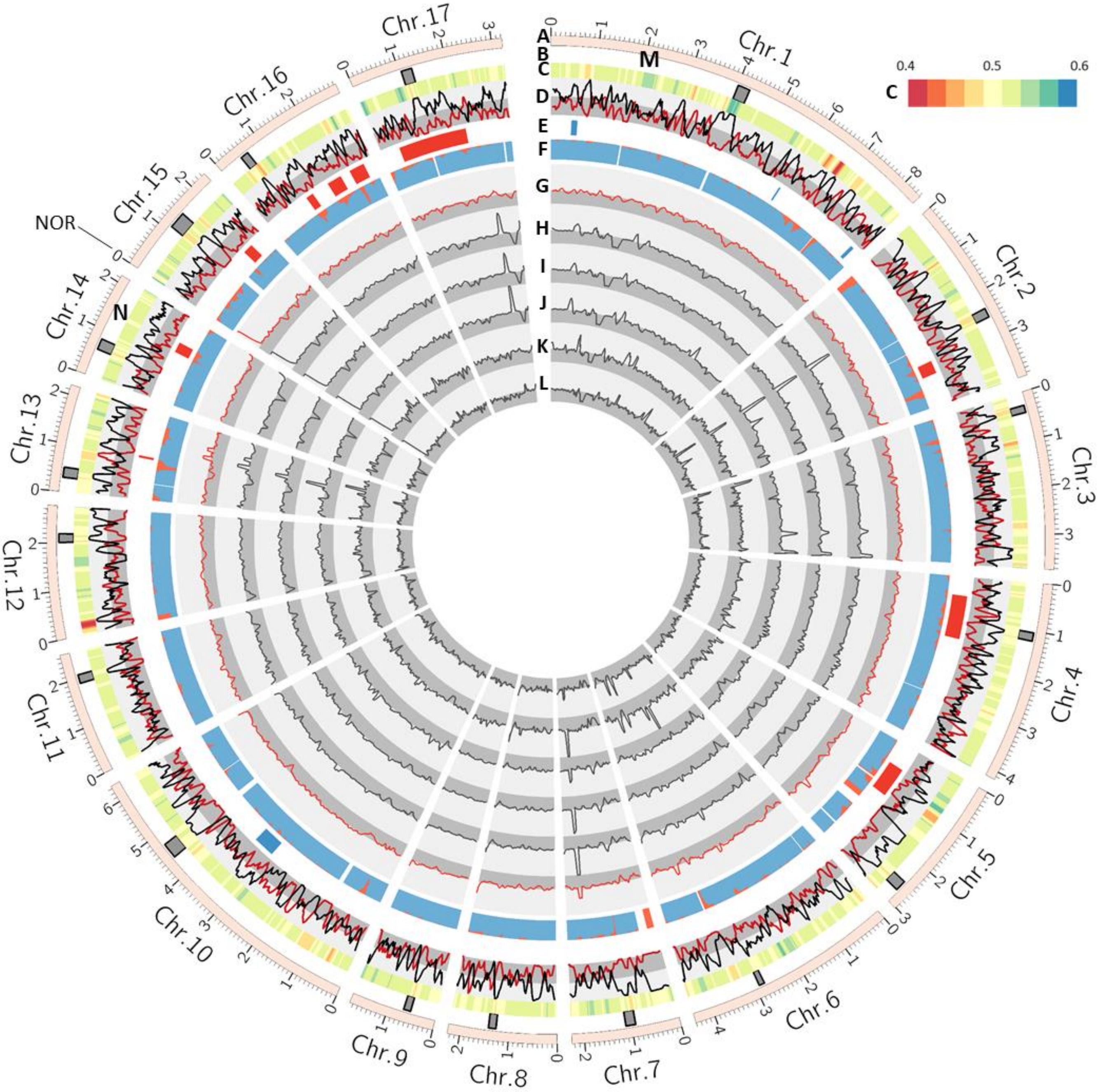


**Supplementary Fig. 14. Read depth across the *P. effusa* UA202013 assembly for six different *P. effusa* isolates.** Tracks A to D are the same a Fig. 1A–D. Track E is Fig. 1F. Track F is Fig. 1G. Tracks G to L are an expansion of Fig. 1I, displaying the normalised read depth of six isolates. Track G is UA202013, H is Pfs12, I is Pfs14, J is R14, K is Pfs13, and L is R13. The background indicated 0x to 1x (dark grey) and 1x to 3x (light grey). All results except UA202013 were generated with Illumina short reads. UA202013 was generated using PacBio HiFi reads. The coverage of UA202013 aligned to itself was uniform, except for sequence around the nucleolus organised region (NOR) on Chr. 15. All isolates had elevated coverage around the NOR. The read coverage profiles of Pfs12, Pfs14, and R14 were similar but not identical. As were the read coverage profiles or Pfs13 and R13. This suggests isolates collected from different studies have similar genotypes. Short read alignments revealed several regions of low coverage, including regions of the genome with high AT content (see M). N) A region on Chr. 14 had low short read coverage and colocalizes with a cluster of effectors. This region may contain an avirulence protein unique to UA202013 compared to the other isolates.

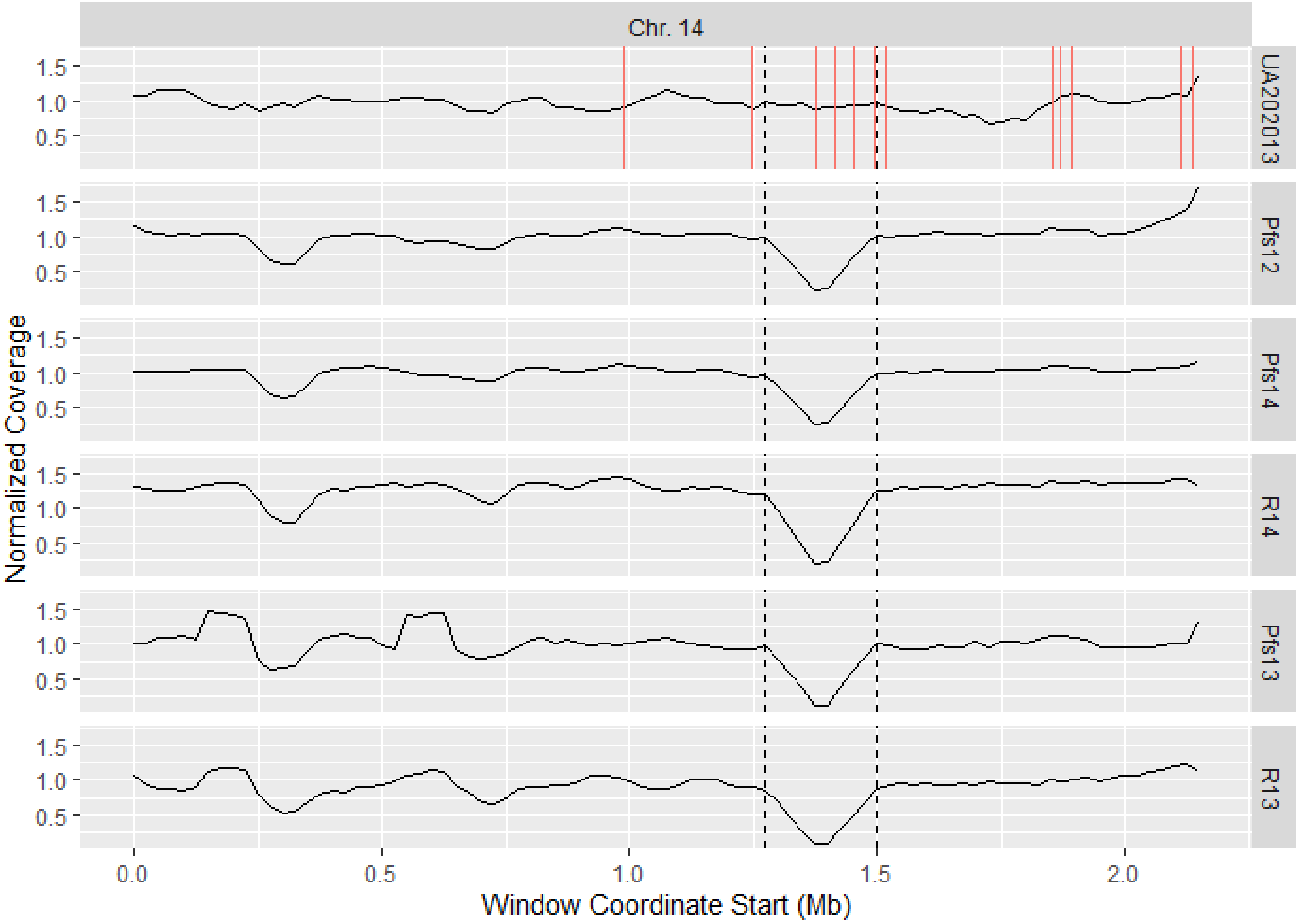


**Supplementary Fig. 15. Read depth across Chr. 14 of the *P. effusa* UA202013 assembly. The six panels represent a different isolate labelled on the right.** The low coverage region identified across all isolates except UA202013, in Supplementary Fig. 14N, is indicated by black lines. Red lines on the UA202013 panel represent the coordinates of effectors annotated in UA202013. The four effectors within the region cluster at high identity to one another.

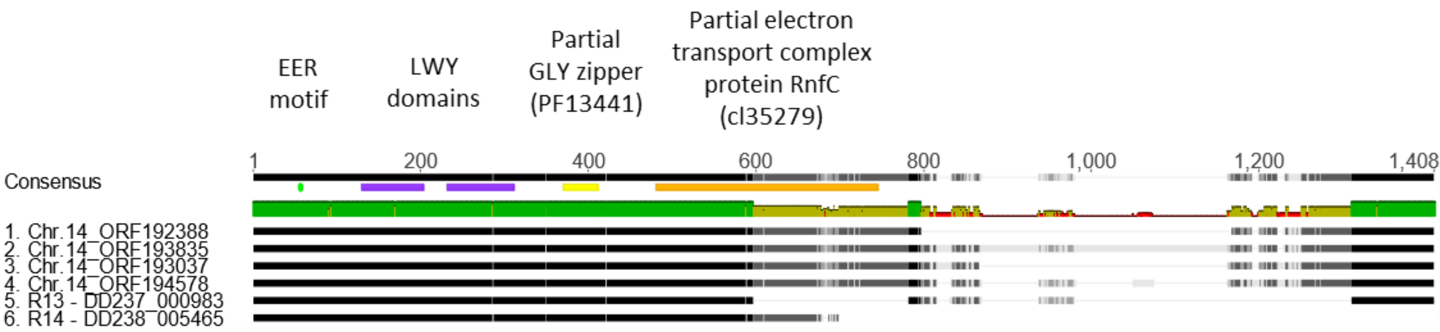


**Supplementary Fig. 16. Alignment of the effectors of *P. effusa* identified in the low coverage window of Chr. 14, plus single homologs annotated in R13 and R14.** The annotated features of the protein include two LWY domains and an EER motif. Surprisingly, two additional, partial domains were also detected using the NCBI conserved domain search. In the R13 and R14 homologous annotations the electron transport complex protein RnfC (cl35279) domain is disrupted. Annotation of UA20213 submitted to NCBI and awaiting the assignment of accession identities. Similar protein sequences can be found by using the NCBI accessions of P. effusa isolates R13 or R14.

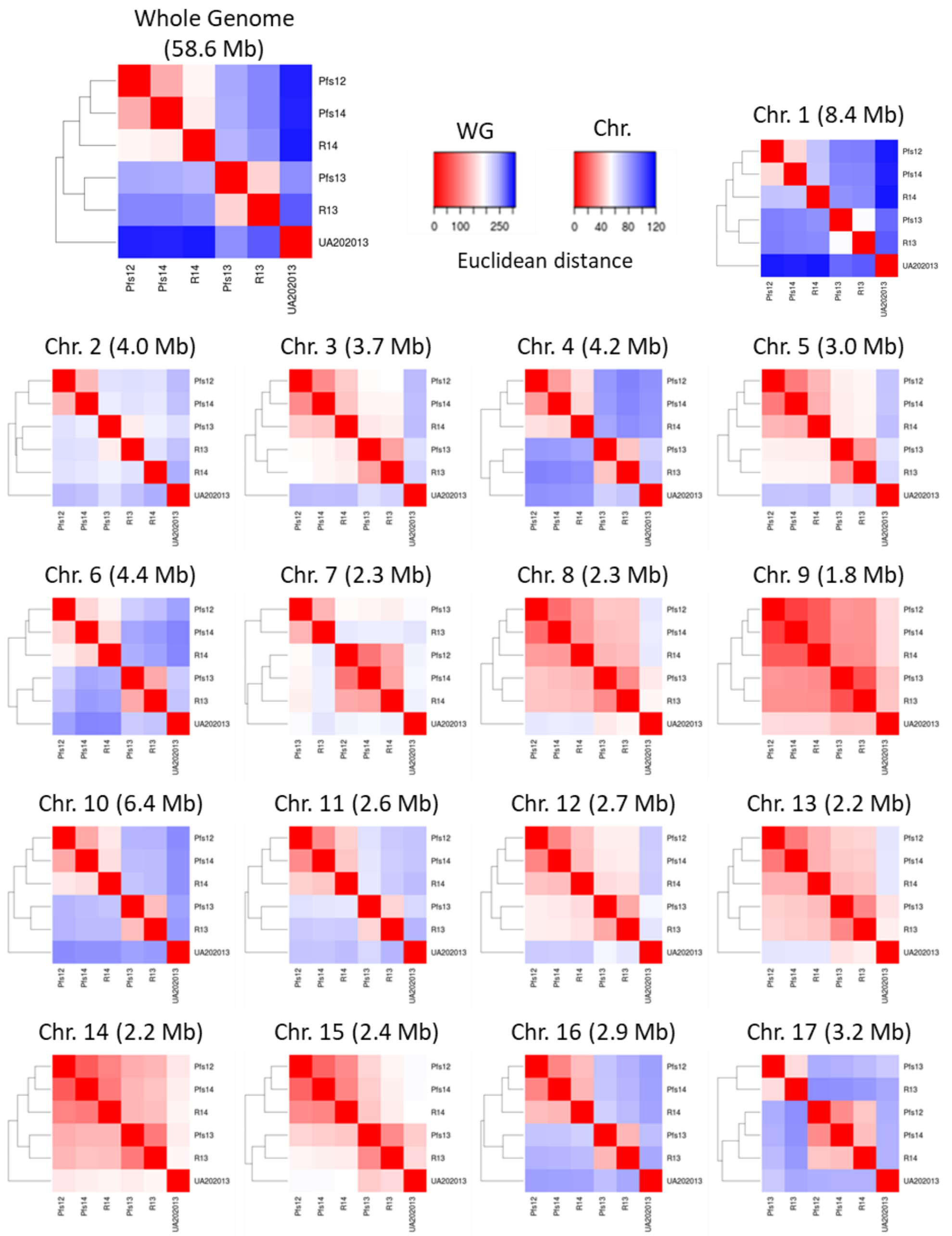


**Supplementary Fig. 17. Clustering of isolates by SNPs called against the *P. effusa* UA202013 assembly.** This analysis considered bi-allelic SNPs called as either reference homozygous, heterozygous, or alternative homozygous. The larger top left heatmap considers all variants called against the whole genome assembly and is on an independent scale (WG). The 17 smaller heatmaps display the calculated Euclidean distances per chromosome and are on a common scale (Chr.). The number of variants identified per chromosome correlated with chromosome length (Pearson's correlation = 0.99). The mean Euclidean distance between isolates correlated with chromosome length (Pearson's correlation = 0.84) due to larger chromosomes having more variants. Therefore, larger chromosomes such as Chr. 1 contain more variants and have a larger mean Euclidean distance than smaller chromosomes such as Chr. 9. *P. effusa* isolates R13 (Fletcher *et al.* 2018) and Pfs13 (Feng *et al.* 2018) represent the same race phenotype and clustered together as pairs. *P. effusa* isolates R14 (Fletcher *et al.* 2018) and Pfs14 (Feng *et al.* 2018) are the same race but did not cluster as pairs; instead, Pfs14 clustered with Pfs12 on every chromosome.

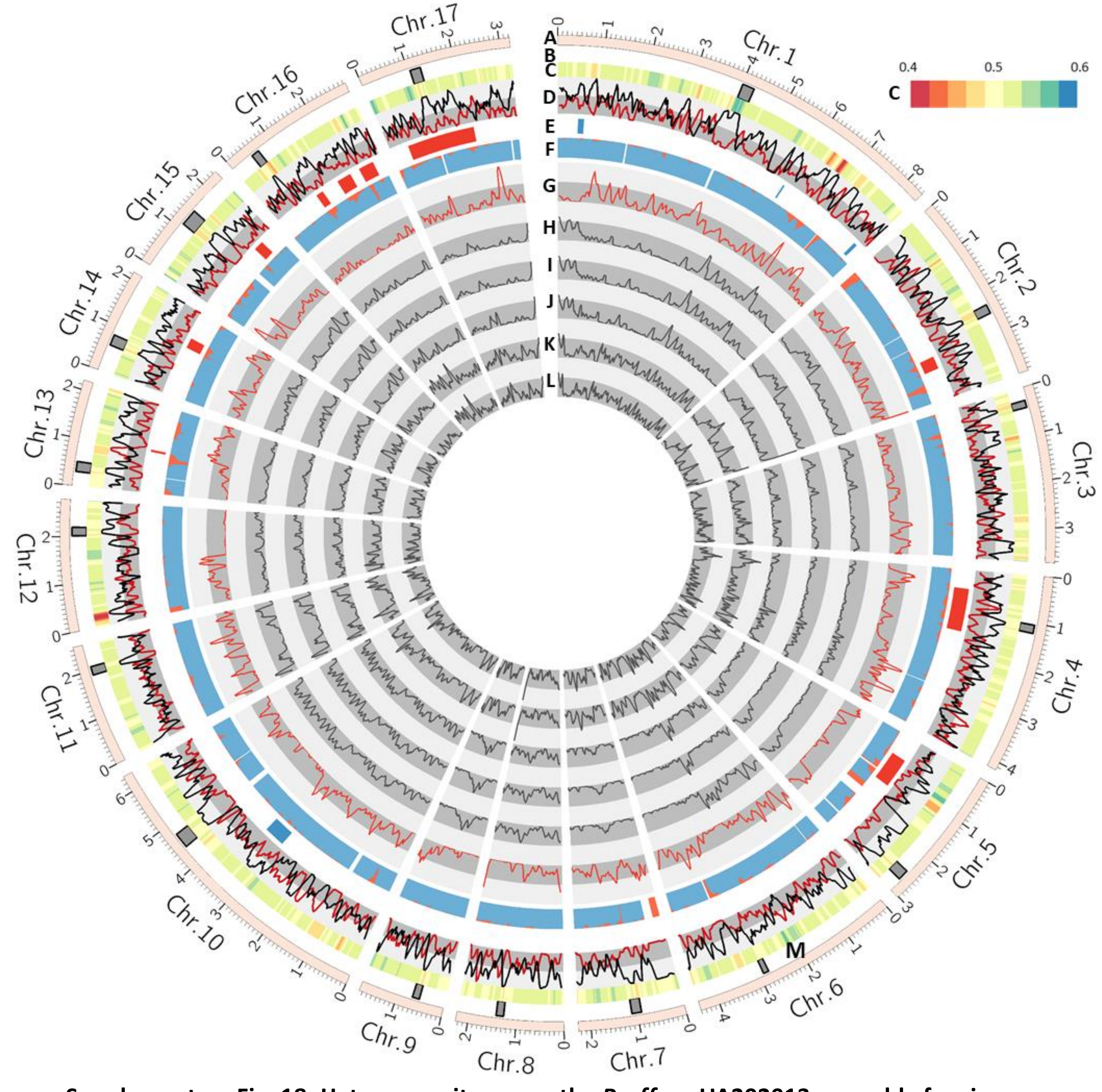


**Supplementary Fig. 18. Heterozygosity across the *P. effusa* UA202013 assembly for six different *P. effusa* isolates.** Tracks A to D are the same a Fig. 1A–D. Track E is Fig. 1F. Track F is Fig. 1G. Tracks G to L are levels of heterozygosity per window for six isolates and were derived from Fig. 1K. Track G is UA202013, H is Pfs12, I is Pfs14, J is R14, K is Pfs13, and L is R13. Results only consider SNPs called as heterozygous, differing from Fig. 1K, which displays heterozygous and homozygous alternative SNPs. As with read-depth (Supplementary Fig. 14), the heterozygosity profiles of Pfs12, Pfs14, and R14 were similar but not identical, as were the heterozygosity profiles of Pfs13 and R13. This suggests isolates collected in different studies have similar genotypes. M) Heterozygosity on Chr. 6 differed between Pfs12 and race 14 isolates Pfs14 and R14. Loss of heterozygosity was previously proposed as the mechanism of adaptation between the race 12 and race 14 phenotype (Lyons *et al.* 2016). In the assembly of UA202013, this region contained effectors but did not overlap a cluster of effectors. A second, smaller region differing in heterozygosity between these isolates was identified on Chr. 14. This region contained no effectors in the assembly of UA202013.

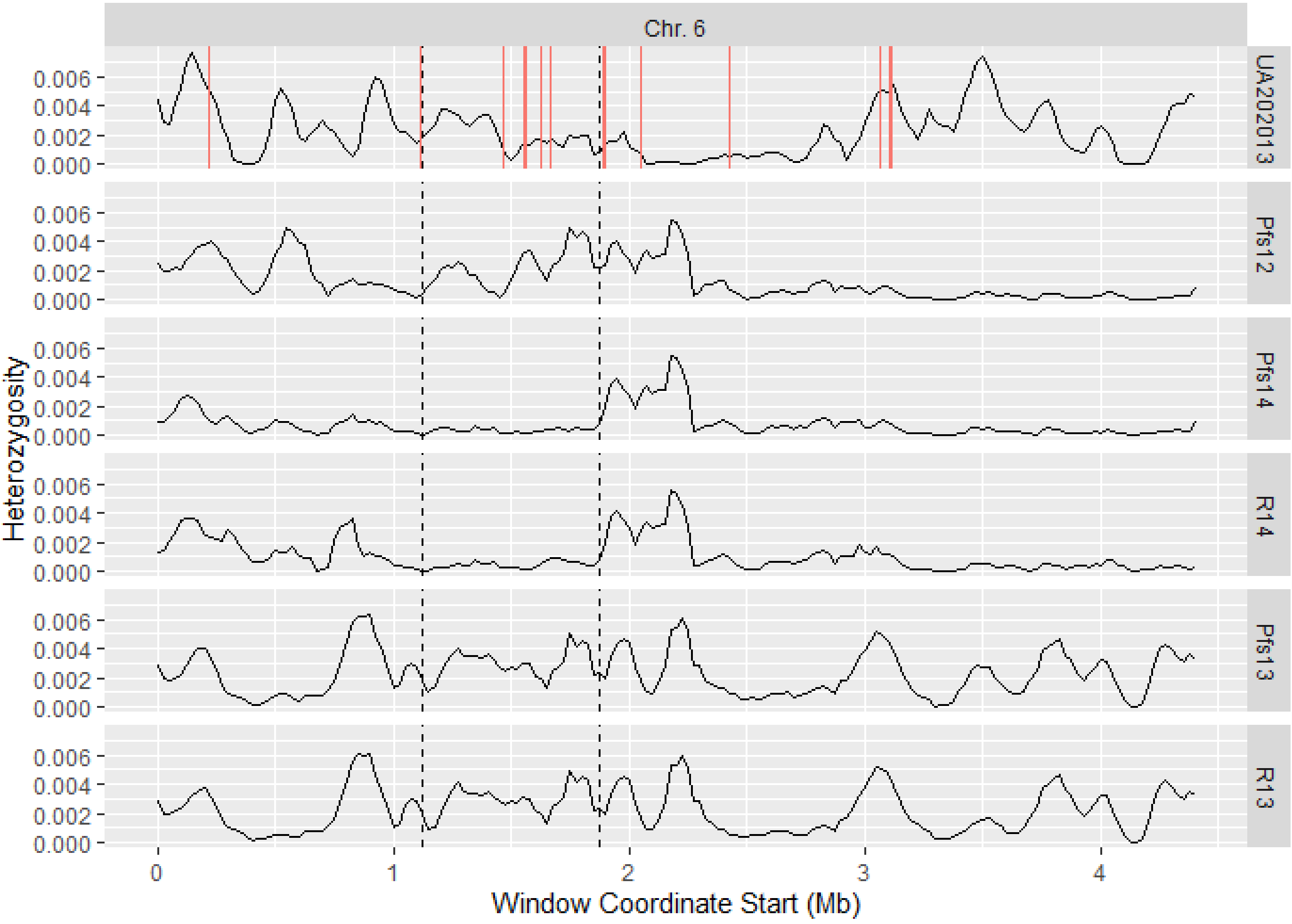


**Supplementary Fig. 19. Heterozygosity across Chr. 6 of the *P. effusa* UA202013 assembly for six different *P. effusa* isolates.** The six panels represent a different isolate labelled to the right. The loss of heterozygosity region identified across race 14 isolates compared to Pfs12, in Supplementary Fig. 18M, is indicated by black lines. Red lines on the UA202013 panel represent the coordinates of effectors annotated in UA202013. The six UA202013 effectors annotated in the region did not form a cluster of effectors. *P. effusa* isolates Pfs12, Pfs14, and R14 appear more homozygous across the entirety of Chr. 6.

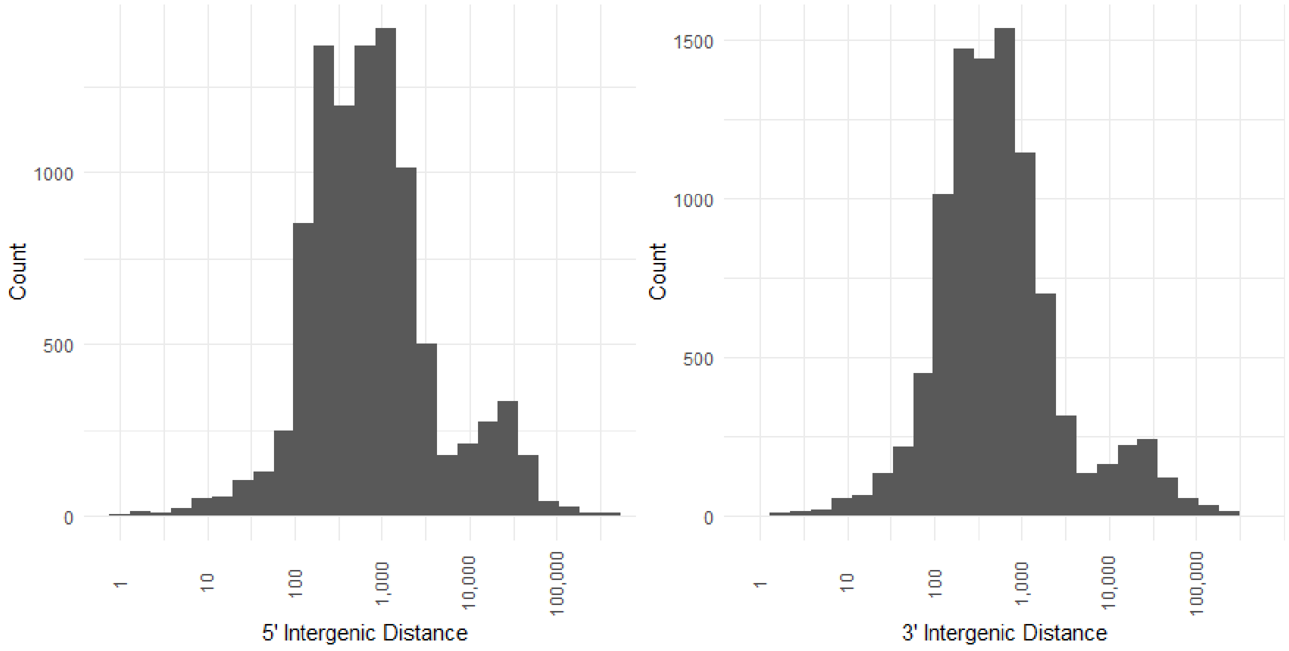


**Supplementary Fig. 20. Intergenic distances between genes were bimodal in *Peronospora effusa*.** This is consistent with the two-speed hypothesis (Dong *et al.* 2015).
