## Supplementary Tables 1 to 2 for "Ancestral chromosomes for the Peronosporaceae inferred from a telomere-to-telomere genome assembly of *Peronospora effusa*"

**Supplementary Table 1. List of oomycete assemblies used for orthology analysis.**

*Achlya hypogyna*
*Albugo candida*
*Albugo laibachii*
*Aphanomyces astaci*
*Aphanomyces invadans*
*Bremia lactucae*
*Globisporangium irregulare*
*Globisporangium ultimum* BR650
*Globisporangium ultimum* DAOMBR144
*Hyaloperonospora arabidopsidis*
*Peronospora effusa* R13
*Peronospora effusa* R14
*Peronospora effusa* UA202013
*Peronospora tabacina* S26
*Peronospora tabacina* J2
*Phytophthora cactorum*
*Phytophthora capsici*
*Phytophthora cinnamomi*
*Phytophthora infestans*
*Phytophthora kernoviae*
*Phytophthora megakarya*
*Plasmopara muralis*
*Phytophthora nicotianae* AM5888
*Phytophthora nicotianae* R0
*Phytophthora palmivora*
*Phytophthora parasitica*
*Phytophthora ramorum*
*Phytophthora sojae*
*Phytopythium vexans*
*Plasmopara halstedii*
*Plasmopara viticola*
*Pythium insidiosum*
*Pythium iwayamai*
*Pythium aphanidermatum*
*Pythium arrhenomanes*
*Saprolegnia diclina*
*Saprolegnia parasitica*
*Thraustotheca clavlata*

**Supplementary Table 2. Normalized read-depth of genes encoding metallophosphatase domain containing proteins used in Fig. 2C.**

| | UA202013 | Pfs12 | Pfs13 | Pfs14 | R13 | R14 |
|---|---|---|---|---|---|---|
| Chr. 1 | 0.79x | 1.17x | 1.3x | 1.1x | 0.85x | 0.99x |
| Chr. 2 | 0.67x | 0.89x | 0.6x | 0.79x | 1.1x | 1.03x |
| Chr. 13 | 1.13x | 0.96x | 0.96x | 0.97x | 0.95x | 0.99x |
| Chr. 14 | 1.4x | 0.97x | 0.69x | 0.95x | 1.12x | 1.07x |
